# Separable neural signatures of confidence during perceptual decisions

**DOI:** 10.1101/2021.04.08.439033

**Authors:** T. Balsdon, P. Mamassian, V. Wyart

**Affiliations:** Laboratoire des Systèmes Perceptifs (CNRS UMR 8248), DEC, ENS, PSL University, 75005 Paris, France; Laboratoire de Neurosciences Cognitives et Computationnelles (Inserm U960), DEC, ENS, PSL University, 75005, Paris, France

**Author notes:** Equal contributors (P. Mamassian, V. Wyart).

## Abstract

Perceptual confidence is an evaluation of the validity of perceptual decisions. While there is behavioural evidence that confidence evaluation differs from perceptual decision-making, disentangling these two processes remains a challenge at the neural level. Here we examined the electrical brain activity of human participants in a protracted perceptual decision-making task where observers tend to commit to perceptual decisions early whilst continuing to monitor sensory evidence for evaluating confidence. Premature decision commitments were revealed by patterns of spectral power overlying motor cortex, followed by an attenuation of the neural representation of perceptual decision evidence. A distinct neural representation was associated with the computation of confidence, with sources localised in the superior parietal and orbitofrontal cortices. In agreement with a dissociation between perception and confidence, these neural resources were recruited even after observers committed to their perceptual decisions, and thus delineate an integral neural circuit for evaluating perceptual decision confidence.

## Introduction

Whilst perception typically feels effortless and automatic, it requires probabilistic inference to resolve the uncertain causes of essentially ambiguous sensory input (Helmholtz, 1856). Human observers are capable of discriminating which perceptual decisions are more likely to be correct using subjective feelings of confidence (Pollack and Decker, 1958). These feelings of perceptual confidence have been associated with metacognitive processes (Fleming and Daw, 2017) that enable self-monitoring for learning (Veenman, Wilhelm, & Beishuizen, 2004) and communication (Bahrami et al., 2012; Frith, 2012). We are only just beginning to uncover the complex functional role of metacognition in human behaviour, and outline the computational and neural processes that enable metacognition. The study of perceptual confidence offers promising insight into metacognition, because one can use our detailed knowledge of perceptual processes to isolate factors which affect the computation of perceptual confidence.

At the computational level, perceptual decisions are described by sequential sampling processes (Vickers, 1970; Ratcliff, 1978), in which noisy samples of evidence are accumulated over time, until there is sufficient evidence to commit to a decision. The most relevant information for evaluating perceptual confidence is the quantity and quality of evidence used to make the perceptual decision (Vickers, 1979; Kepecs et al., 2008; Moreno-Bote, 2010). At the neural level, perceptual confidence could therefore follow a strictly serial circuit: Relying only on information computed by perceptual processes, with any additional processes contributing only to transform this information for building the confidence response required by the task. Indeed, confidence (or a non-human primate proxy for confidence) can be reliably predicted from the firing rates of neurons coding the perceptual decision itself (Kiani and Shadlen, 2009), suggesting that confidence may be a direct by-product of perceptual processing. However, a large body of behavioural studies suggest that the computation of confidence is not strictly serial. Confidence can integrate additional evidence after the observer commits to their perceptual decision (Baranski and Petrusic, 1994; Pleskac and Busemeyer, 2010), and while this continued evidence accumulation could incorporate only perceptual information, it implies that confidence evaluation does not directly follow from perceptual decision commitment (and therefore involves at least partially dissociable neural processes).

There is also evidence that perceptual confidence can rely on separate (non-perceptual) sources of information, such as decision time (Kiani, Corthell, and Shadlen, 2014) and attentional cues (Denison et al., 2018). This suggests that the processes involved in the computation of perceptual confidence may not be reduced to the same processes as for the perceptual decision. Higher-order theories of metacognition propose a framework in which specialised metacognitive resources could be recruited for computing confidence across all forms of decision-making (a general metacognitive mechanism). Indeed, there is some evidence that confidence precision is correlated across different cognitive tasks (such as memory and perception; Mazancieux et al., 2018), suggesting a common source of noise affecting the computation of confidence across tasks (on top of the sensory noise; Bang, Shekhar, and Rahnev, 2019; Shekhar and Rahnev, 2020).

It is reasonable to expect that a general metacognitive mechanism relies on processing in higher order brain regions. Several experiments have linked modulations in confidence with activity in a variety of subregions of the prefrontal cortex (including the orbitofrontal cortex, Masset et al., 2020, Lak et al., 2014; right frontopolar cortex, Yokoyama et al., 2010; rostro-lateral prefrontal cortex, Fleming et al., 2012, Geurts et al., 2021; inferior frontal sulcus, medial frontal sulcus and medial frontal gyrus, Cortese et al., 2016; see also Vaccaro and Fleming, 2018, for a meta-analysis). Moreover, disrupting the processing in subregions of the prefrontal cortex (Rounis et al., 2010; Lak et al., 2014; Fleming et al., 2014) tends to impair (though not obliterate) the ability to appropriately adjust behavioural confidence responses, whilst leaving perceptual decision accuracy largely unaffected (though these results can be difficult to replicate, Bor et al., 2017; Lapate et al., 2020, and may not generalise to metacognition for memory; Fleming et al., 2014). A challenge in this literature is in specifically relating the neural processing to the computation of confidence, as opposed to transforming confidence into a behavioural response, or a downstream effect of confidence, such as the positive valence (and sometimes reward expectation) accompanying correct decisions. Moreover, identifying how these neural mechanisms could be separable from the underlying perceptual processes is important for understanding the computational architecture of metacognition.

One promising avenue of research for separating the mechanisms of metacognition from perceptual processes has been to utilise tasks where the observer may integrate additional evidence for confidence after they have committed to their perceptual decision (Murphy et al., 2015; Fleming et al., 2018), which presumably relies on processing independent of the perceptual decision. These studies show that post-decisional changes in confidence magnitude correlate with signals from the posterior medial frontal cortex. However, these signals could reflect processes occurring downstream of confidence, such as an emotional response to the error signal, which has been shown to drive medial frontal activity more strongly than decision accuracy (Gehring and Willoughby, 2002). Further research is therefore required to link neural processes specifically with the computation of perceptual confidence.

In this experiment we aim to identify the neural processes specifically contributing to the computation of confidence, in a paradigm in which these processes can be delineated from those of perceptual decision-making. We exploit a protracted decision-making task in which the evidence presented to the observer can be carefully controlled. On each trial, the observer is presented with a sequence of visual stimuli, oriented Gabor patches, which offer a specific amount of evidence towards the perceptual decision. The orientations are sampled from one of two overlapping circular Gaussian distributions, and the observer is asked to categorise which distribution the orientations were sampled from. We manipulate the amount of evidence presented such that the observer tends to covertly commit to their perceptual decision before evidence presentation has finished, whilst continuing to monitor ongoing evidence for assessing their confidence (Balsdon et al., 2020). These covert decisions are evident from behaviour and computational modelling, and we show similarities between the neural processes of decision-making across conditions of immediate and delayed response execution.

To examine the computation of confidence, we compare human behaviour to an optimal observer who perfectly accumulates all the presented evidence for perceptual decisions and confidence evaluation. The optimal observer must accurately encode the stimulus orientation, the decision update relevant for the categorisation, and add this to the accumulated evidence for making the perceptual decision. We uncover dynamic neural representations of these variables using model-based electroencephalography (EEG), and examine how the precision of these representations fluctuate with behavioural precision. We find two distinct representations of the accumulated evidence. The first one reflects the internal evidence used to make perceptual decisions. The second representation reflects the internal evidence used to make confidence evaluations (separably from the perceptual evidence), and is localised to the superior parietal and orbitofrontal cortices. Whilst the perceptual representation is attenuated following covert decisions, the confidence representation continues to reflect evidence accumulation. This is consistent with a neural circuit that can be recruited for confidence evaluation independently of perceptual processes, providing empirical evidence for the theoretical dissociation between perception and confidence.

## Results

### Preview

We present analyses to address two key hypotheses in this experiment: First, that observers are prematurely committing to their perceptual decisions whilst continuing to monitor additional evidence for evaluating their confidence. And second, that there are separable neural signatures of the evaluation of confidence during perceptual decision-making. To address the first hypothesis, we use a combination of behavioural analyses and computational modelling, and in addition, show that the EEG signatures of response preparation are triggered from the time of decision commitment, even when this occurs seconds prior to the response cue. To address the second hypothesis, we use the stimulus evoked responses in EEG to trace the representation of the presented evidence throughout each trial. We show that these neural representations of the optimal accumulated decision evidence are less precise when the observers’ behavioural responses were also less precise relative to optimal. We use this to isolate clusters of activity that specifically reflect the internal evidence used for observers’ confidence evaluations beyond the presented evidence. We then localise the sources of this activity, and relate these processes back to observers’ eventual confidence ratings.

### The computational architecture of perceptual confidence

Human observers (N = 20) performed two versions of the task whilst EEG was recorded. Across the two tasks, 100 predefined sequences of oriented Gabors were repeated for each observer, with stimuli presented as described in **Figure 1a**. In the Free task, the sequence continued until observers entered their perceptual decision (**Figure 1b**), indicating which category (**Figure 1d**) they thought the orientations were sampled from. Observers were instructed to enter their response as soon as they ‘felt ready’, on three repeats of each predefined sequence (300 trials in total). In the Replay task (**Figure 1c**), observers were shown a specific number of samples and could only enter their response after the response cue. After entering their perceptual decision, they made a confidence evaluation, how confident they were that their perceptual decision was correct, on a 4-point scale. Importantly, the number of samples shown in the Replay task was manipulated relative to the Free task, in three intermixed conditions: in the Less condition, they were shown two fewer than the minimum they had chosen to respond to over the three repeats of that predefined sequence in the Free task; in the Same condition they were shown the median number of samples; and in the More condition, four more than the maximum. The variability across repeats in the Free task means that in the More condition, observers were show at least four additional stimuli, but often more than that. There is an optimal way to perform this task, in the sense of maximising perceptual decision accuracy across trials. The optimal computation takes as decision evidence the log probability of each orientation given the category distributions (**Figure 1d**) and accumulates the difference in this evidence for each category (**Figure 1e**, Drugowitsch et al., 2016). We refer to the accumulated difference in log probabilities as the optimal presented evidence, *L*. Human observers may have a suboptimal representation of this evidence, *L\**, and we estimate the contribution of different types of suboptimalites (specifically, inference noise, and a temporal integration bias) with the help of a computational model (full details in **Methods** and **Supplementary Note 1**).

**Figure 1.**
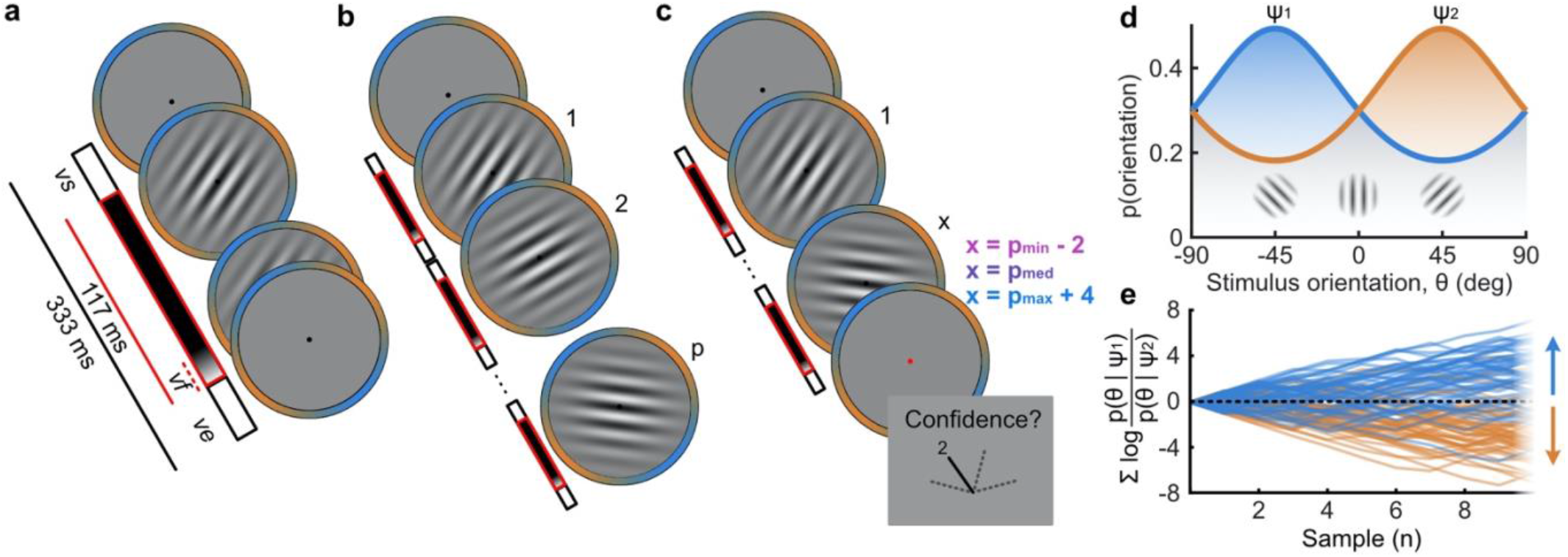
Procedure. **a)** Stimulus presentation: stimuli were presented at an average rate of 3 Hz, but with variable onset and offset (**vs** ∈ **[83, 133]** ms, **vs**_**s**_ + **ve**_**s–1**_ ≥ **216** ms; see **Methods**). Stimuli were presented within a circular annulus which acted as a colour guide for the category distributions. The colour guide and the fixation point were present throughout the trial. **b)** Free task: on each trial observers were presented with a sequence of oriented Gabors, which continued until the observer entered their response (or 40 samples were shown). 100 sequences were predefined and repeated three times. **c)** Replay task: The observer was presented with a specific number of samples and could only enter their response after the cue (fixation changing to red). The number of samples (x) was determined relative to the number the observer chose to respond to on that same sequence in the Free task (p). There were three intermixed conditions, Less (x = p_min_ – 2; where p_min_ is the minimum p of the three repeats), Same (x = p_med_; where p_med_ is the median p) and More (x = p_max_ + 4; where p_max_ is the maximum p of the three repeats of that predefined sequence). **d)** Categories were defined by circular Gaussian distributions over the orientations, with means −45° (**ψ**_**1**_, blue) and 45° (**ψ**_**2**_, orange), and concentration **κ** = **0.5**. The distributions overlapped such that an orientation of 45**°** was most likely drawn from the orange distribution but could also be drawn from the blue distribution with lower likelihood. **e)** The optimal observer accumulates the difference in the evidence for each category, which is defined as the log probability of the sample orientation (**θ**) given the distributions. The perceptual decision is determined by the sign of the accumulated evidence, where the evidence accumulated across more samples better differentiates the true categories (example evidence traces are coloured by the true category).

Based on our previous findings (Balsdon et al., 2020) we expected observers to prematurely commit to perceptual decisions in the More condition, whilst continuing to monitor sensory evidence for evaluating their confidence. Replicating these previous results (Balsdon et al., 2020), we found that perceptual decision sensitivity (d’) was significantly decreased with just two fewer stimuli in the Less condition compared to those same (*p*_*min*_) trials in the Free task (Wilcoxon sign rank *Z* = 3.88, *p* < 0.001, Bonferroni corrected for three comparisons, **Figure 1a**), but four additional stimuli (**Figure 1b**) in the More condition resulted in only a small but not significant increase compared to the *p*_*max*_ trials in the Free task (*Z* = −1.53, *p* = 0.13, uncorrected). There was also no significant difference for the Same condition (*Z* = 1.21, *p* = 0.23, uncorrected; **Figure 2a**).

**Figure 2.**
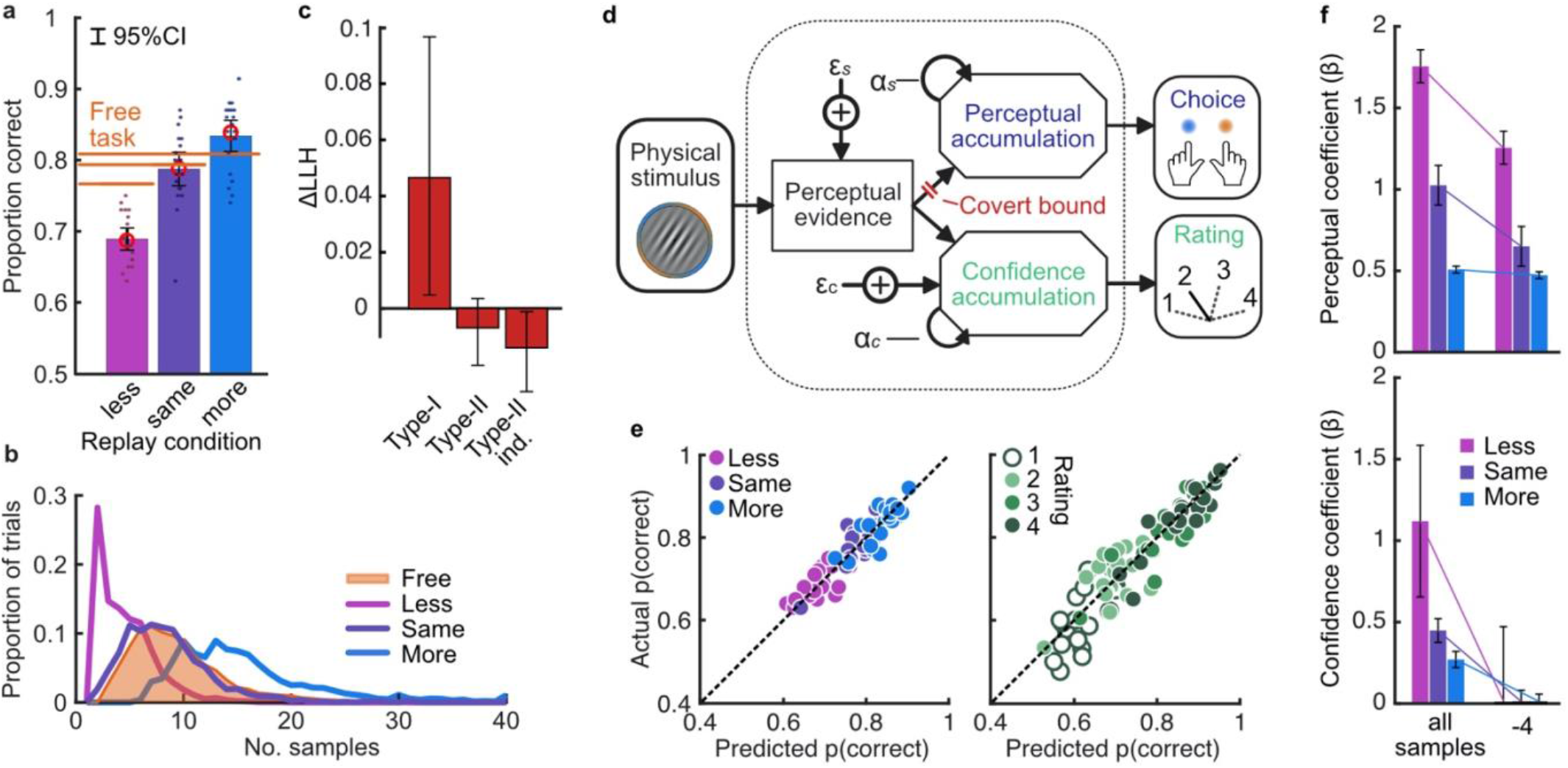
Behaviour and computational modelling. **a)** Proportion correct in each condition of the Replay task, relative to the Free task (orange horizontal lines). Individual data are shown in scattered points, error bars show 95% between-(thin) and 95% within-(thick) subject confidence intervals. Open red markers show the model prediction. **b)** Distributions of the number of samples per trial in the Free task, and Replay task conditions (over all observers).**c)** Difference in log-likelihood of the models utilising a covert bound relative to the models with no covert bound. On the left, the model fitting perceptual decisions only. The middle bar shows the difference in log-likelihood of the fit to confidence ratings with identical perceptual and confidence bounds. The right bar shows the difference in log-likelihood of the fit to confidence ratings of the model with an independent bound for confidence evidence accumulation. Error bars show 95% between-subject confidence intervals. **d)** The computational architecture of perceptual and confidence decisions, based on model comparison. Perceptual and confidence decisions accumulate the same noisy perceptual evidence, but confidence is affected by additional noise (ε_c_) and a separate temporal bias (α_c_) This partial dissociation allows Type-II accumulation to continue after the observer has committed to a perceptual decision. **e)** Predicted proportion correct compared to actual proportion correct for each observer, based on the fitted model parameters of the final computational model. The left panel shows proportion correct split by condition, and the right, split by confidence rating. **f)** Regression coefficients from the GLM analysis showing the relationship between the optimal evidence L, and observers’ perceptual (top) and confidence (bottom) responses for trials split by condition. The right set of bars show the same analysis but with evidence accumulated up to four samples from the response cue.

This lack of substantial increase in performance in the More condition could be the result of either a performance ceiling effect or a premature commitment to the perceptual decision. The former explanation reflects a limitation of the perceptual evidence accumulation process, whereas the latter refers to an active mechanism that ignores the final sensory evidence. We compared these two hypotheses using a computational modelling approach (Balsdon et al., 2020; see **Methods**). Specifically, we compared a model in which performance in the More condition is limited by the suboptimalities evident from the Same and the Less conditions (inference noise, and temporal integration bias, see **Methods** and **Supplementary Note 1**), to a model in which performance could be impacted by a covert bound at which point observers commit to a decision irrespective of additional evidence. Cross-validated model comparison provided significant evidence that observers were implementing a covert bound (mean relative increase in model log-likelihood = 0.048, bootstrapped *p* = 0.001, **Figure 2c**). The winning model provided a good description of the data (red open markers in **Figure 2a**, and individual participants in **Figure 2e**).

In contrast to what we found for the perceptual decision, there was no evidence that observers were implementing a covert bound on confidence: Implementing the same bound as the perceptual decision did not improve the fit (relative improvement with bound = −0.007, bootstrapped *p* = 0.11, uncorrected) and an independent bound actually significantly *reduced* the fit compared to continued accumulation (relative improvement = −0.014, *p* = 0.022, Bonferroni corrected for two comparisons; **Figure 2c**). We obtained further distinctions between perceptual and confidence processes through computational modelling: additional noise was required to explain the confidence ratings, along with a separate temporal bias. The best description of both perceptual and confidence responses was provided by a partially dissociated computational architecture (full details in **Supplementary Note 1**), where perceptual and confidence decisions are based on the same noisy representation of the sensory evidence, but confidence accumulation incurs additional noise and can continue after the completion of perceptual decision processes (**Figure 2d**, and the predictions of this model for individual participants are show in **Figure 2e**). These computational differences between perceptual decisions and confidence evaluations suggest deviations between the internal evidence on which observers base their perceptual and confidence decisions (see **Supplementary Note 2** for model simulations).

These modelling results are supported by an analysis using general linear models to examine the relationship between the optimal presented evidence, *L*, and observers’ behaviour in the perceptual decision and confidence evaluation. As stated above, *L* is the evidence that which maximises the probability of a correct response: the accumulated difference in the log probabilities of the presented orientations given the category distribution (**Figure 1e**). First, we find the presented evidence accumulated over all samples does explain substantial variance in observers’ perceptual decisions (average *β* = 0.77, *t*(19) = 6.48, *p* < 0.001), and confidence evaluations (with the evidence signed by the perceptual response; *β* = 0.24, *t*(19) = 6.46, *p* < 0.001). This suggests that the internal evidence that observers were using to make their responses, *L\**, correlated significantly with the optimal evidence *L* (as has been found previously; Drugowitsch et al., 2016). Second, the total accumulated evidence in the More condition was not a significantly better predictor of the observers’ perceptual decisions than the evidence up to four samples prior to the response (average difference in *β* = 0.034, *t*(19) = 1.63, *p* = 0.12), while for the Same and Less conditions the total accumulated evidence was a significantly better predictor (Less: *t*(19) = 4.99, *p* < 0.001; Same: *t*(19) = 3.11, *p* = 0.006; causing a significant interaction between condition and sample accumulated to *F*(2,38) = 10.348, *p* = 0.001, Bonferroni corrected for three comparisons, **Figure 2f**, top). This supports the finding from model comparison and behaviour that observers implemented a covert bound on perceptual evidence accumulation. And finally, this interaction was not present when examining how the presented evidence affected confidence evaluations (*F*(2,38) = 3.124, *p* = 0.09, uncorrected, **Figure 2f**, bottom). Rather, the accumulated evidence up to the final sample in the More condition was a significantly better predictor of confidence than the evidence accumulated to four samples from the response (average difference in *β* = 0.26, *t*(19) = 5.33, *p* < 0.001), supporting the prediction from the computational model analysis that observers integrated all the presented evidence for evaluating confidence.

### EEG signatures of premature perceptual decision commitment

The analysis of behaviour and computational modelling so far has suggested that observers were committing to their perceptual decisions early in the More condition and ignoring the additional evidence for their perceptual decision. We questioned the extent of this covert decision commitment, that is, whether observers were going as far as to plan their motor response before the response cue. We examined the neural signatures of the planning and execution of motor responses using a linear discriminant analysis of the spectral power of band-limited EEG oscillations (see **Methods**). Initial analysis suggested the spectral power in the 8 to 32 Hz frequency range (the ‘alpha’ and ‘beta’ bands) could be used to classify perceptual decisions based on lateralised differences over motor cortex (**Supplementary Note 5**). A classifier was trained to discriminate observers’ perceptual decisions at each time-point in a four second window around the response in the Free task (3 seconds prior to 1 second after). This classifier was then tested across time in each condition of the Replay task, to trace the progression of perceptual decision-making in comparison to the Free task (where decisions are directly followed by response execution). If covert decisions lead to early motor response preparation, we would expect asymmetries in cross-classification performance on trials where the observer was likely to have covertly committed to a decision (in the More condition) compared to those trials in which they were unlikely to have committed to their decision (in the Less condition). Indeed, there were opposite asymmetries in the cross-classification of the Less and the More conditions (**Figure 3a**). Statistical comparison revealed substantial clusters of significant differences (**Figure 3b**): Training around −0.78 to 0.44 s from the time of the response in the Free task led to significantly better accuracy testing in the More condition than in the Less condition, prior to when the response was entered (for the cluster testing at −2.5 to −1.6 s *Z_ave_* = 2.04, *p_cluster_* = 0.002; testing at −1.5 to −1 s, *Z_ave_* = 1.95, *p_cluster_* = 0.01; testing at −0.8 to −0.3, *Z_ave_* = 2.32, *p_cluster_* < 0.001). This pattern of findings suggests that observers were not only committing to their perceptual decision early, but already preparing their motor response.

**Figure 3.**
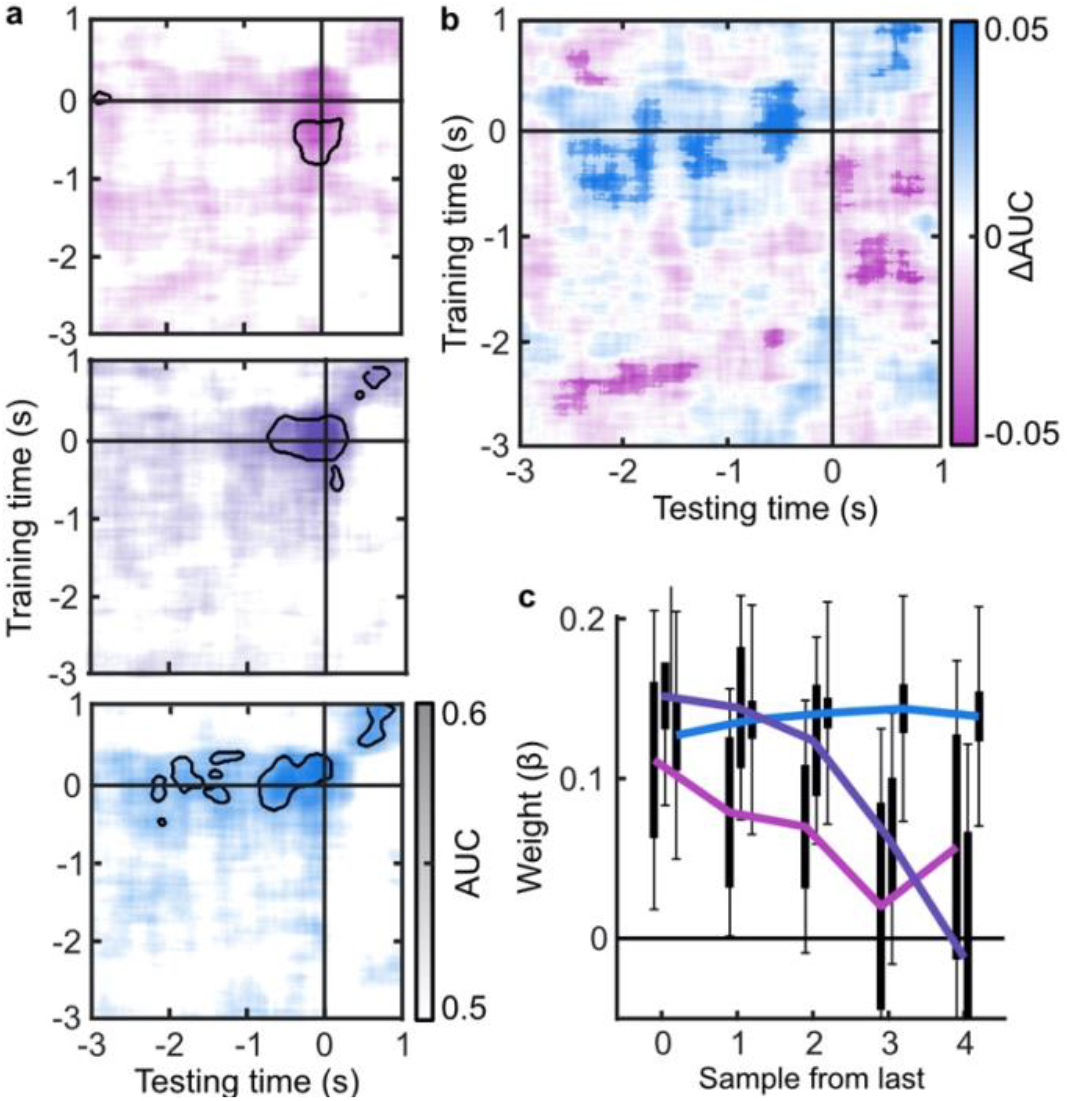
EEG signatures of premature perceptual decisions. **a)** Classifier AUC training at each time-point in the Free task and testing across time in the Less (top), Same (middle), and More (bottom) conditions of the Replay task. Black contours encircle regions where the mean is 3.1 standard deviations from chance (0.5; 99% confidence). **b)** Difference in AUC between the More and Less conditions. Cluster corrected significant differences are highlighted. **c)** The relationship between the evidence accumulated up to n samples prior to the response cue and the strength of the neural signature of response execution in each condition. Error bars show 95% within-(thick) and between-subject (thin) confidence intervals.

As an exploratory analysis, we took the strength of the classifier prediction trained and tested at the time of the response as a trial-wise measure of the decision variable used by the participant to enter a response. We reasoned that the amount of evidence in favour of the decision could influence the assiduity with which observers enter their response. We found that the optimal evidence *L*, accumulated over all samples, could predict the strength of the classifier prediction at response time (mean *β* = 0.11, *t*(19) = 3.89, *p* < 0.001; **Figure 3c**). For the Same and Less conditions, the weight on the accumulated evidence appeared to decrease as evidence was accumulated to samples further prior from the response. But, in the More condition, the evidence accumulated up to four samples prior to the response still predicted the strength of the classifier prediction (*t*(19) = 3.81, *p* = 0.001). This difference between conditions over samples is evidenced by a significant interaction based on a repeated measures ANOVA (*F*(8,152) = 2.429, *p* = 0.05, after Bonferroni correction for three comparisons). Leading up to the response, the accumulated evidence becomes increasingly predictive of the strength of the classifier prediction, except in the More condition, where this prediction is already accurate up to four samples prior to the response: After committing to a perceptual decision, the observer’s perceptual response is no longer influenced by additional evidence.

### Representations of decision evidence in EEG signals

Our main goal was to isolate the neural signatures of the computation of confidence. Observers’ behaviour varied with the optimal evidence *L* presented to them, but the internal evidence on which they based their perceptual decisions and confidence evaluations, *L\**, clearly deviated from *L.* In other words, the observers’ behavioural performance was not optimal. To identify the neural computations underlying human behaviour, we therefore began by isolating the neural signals which correlate with *L*. We then isolated where and when deviations in the neural representation of *L* covary with deviations in *L\** - the internal evidence reflected in observers’ behaviour.

To perform this task the optimal observer must encode the orientation of the stimulus, estimate the decision update based on the categories, and add this to the accumulated evidence for discriminating between the categories (Wyart et al., 2012; Wyart et al., 2015). We examined the neural representation of these optimal variables using a regression analysis with the EEG signals (evoked response, bandpass filtered between 1 and 8 Hz, see **Methods**). At each time point, we used the relationship between the pattern of neural activity and the encoding variables on 90% of the data to predict the encoding variables on the remaining 10% of the data (10-fold cross validation). The precision of the neural representation was calculated as the correlation between the predicted encoding variable and actual encoding variable in the held-out data, across all 10 folds (see **Methods**). **Figure 4a** shows the time course of the precision of the neural representation of stimulus orientation, momentary decision update, and accumulated evidence (*L*), locked to stimulus onset. The precision of the representations of these variables showed distinct time courses and relied on distinct patterns of EEG activity over scalp topography (**Figure 4b**). There was a transient representation of stimulus orientation localised over occipital electrodes. The representation of the momentary decision update was maintained for a longer duration, initially supported by occipital electrodes, then increasingly localised over central-parietal electrodes. The representation of the accumulated evidence was sustained even longer and relied on both frontal and occipital electrodes.

**Figure 4.**
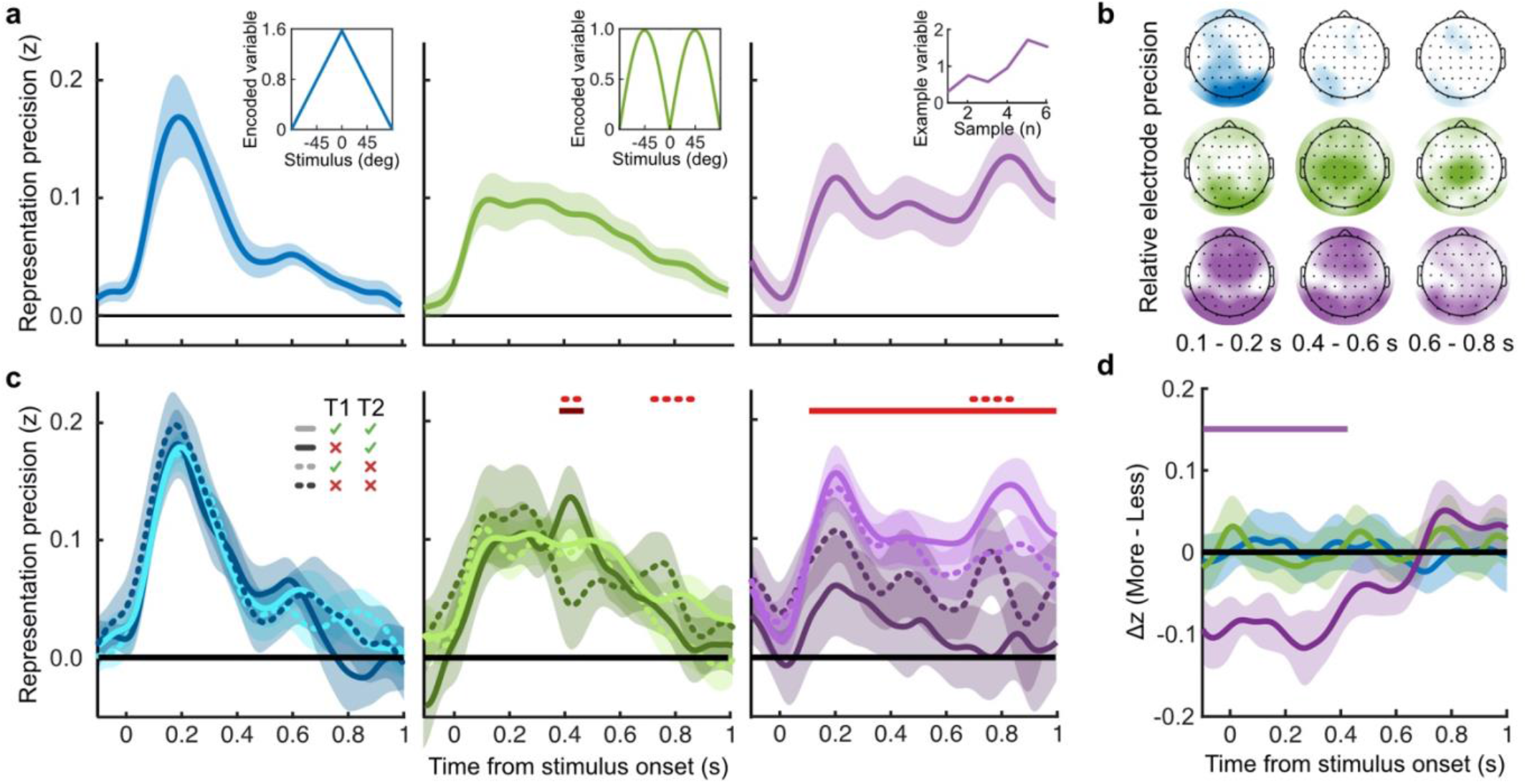
Representation of decision variables. **a)** Representation precision (Fischer transformed correlation coefficient, z) of stimulus orientation (blue, left), momentary decision update (green, middle), and accumulated decision evidence (purple, right). The encoded variables are shown in the insets (the accumulated evidence is the cumulative sum of the momentary evidence signed by the response, only one example sequence is shown). Shaded regions show 95% between-subject confidence intervals. **b)** Relative electrode representation precision over three characteristic time windows (100 – 200 ms, left; 400 – 600 ms, middle; and 600 – 800 ms, right). **c)** Representation precision for epochs leading to optimal and suboptimal perceptual (T1) and confidence (T2) responses. Lighter lines show perceptual decisions that match the optimal response, dashed lines show suboptimal confidence ratings. Dashed red horizontal lines show significant interactions between perceptual and confidence suboptimality. The light red horizontal line shows the significant effect of suboptimal perception and the dark red horizontal line shows the significant effect of suboptimal confidence. Shaded regions show 95% within-subject confidence intervals. **d)** Difference in decoding precision between the More and the Less conditions for epochs corresponding to the last four samples of the trial. The purple horizontal line shows the significant difference in decoding of accumulated evidence.

The internal evidence on which observers base their response, *L\**, can differ from the optimal evidence, *L.* When the eventual behavioural response differs from that predicted by *L*, *L\** is likely to be more different from L. A neural representation of *L* that reflects *L\** (that is, reflecting the underlying processing responsible for behaviour) should also be less precise for samples in these trials. For each variable, we estimated the representation precision separately for epochs leading to behavioural responses that differed from the optimal response (based on *L*), and responses that matched those of the optimal observer (Replay task epochs only; **Figure 4c**; **Supplementary Note 3**). For perceptual decisions, the optimal observer responds with the correct category. For confidence evaluations, the optimal observer gives high confidence on trials with greater than the median evidence (over all trials) for their perceptual response. The precision of the representation of stimulus orientation did not significantly vary based on whether behaviour matched the optimal response. The representation precision of the momentary decision update showed a significant effect for the perceptual decision from 380 to 468 ms (*F_avg_*(1,19) = 7.97, *p_cluster_* = 0.008) and a significant interaction between perceptual and confidence responses from 396 to 468 ms (*F_avg_*(1,19) = 6.66, *p_cluster_* = 0.022) and from 716 to 856 ms (*F_avg_*(1,19) = 10.75, *p_cluster_* < 0.001). The largest effects were seen in the representation precision of the accumulated evidence. Representation precision was significantly reduced in epochs leading to non-optimal perceptual decisions from 108 ms post stimulus onset to the end of the epoch (*F_avg_*(1,19) = 13.65, *p_cluster_* <0.001). In addition, there was a significant interaction with confidence from 696 to 836 ms (*F_avg_*(1,19) = 8.72, *p_cluster_* = 0.005). The precision of the EEG representations therefore showed distinct associations with behaviour.

The presence of a covert bound implies that, after the observer commits to a decision, they no longer incorporate additional evidence for that decision. We should therefore see significant decreases in the precision of representations that specifically contribute to perceptual evidence accumulation. Indeed, the precision of the early representation of accumulated evidence was significantly attenuated for the last four samples of the More condition (in which observers were likely to have already committed to a decision), compared to the last four samples of the Less condition (where observers were unlikely to have committed to a decision; from the start of the epoch to 424 ms, **Figure 4d**; *t_avg_*(19) = −5.19, *p_cluster_*<0.001). These differences in representation precision were not present for the encoding of stimulus orientation, nor the decision update, suggesting that these processes may reflect input to perceptual evidence accumulation, but not the accumulation process itself. As a control analysis, this decreased precision was not evident in a comparison of the first four samples (**Supplementary Note 6**), suggesting this effect on the representation of accumulated evidence is specific to those samples likely to have occurred after perceptual decision commitment, as opposed to those samples in More condition trials per se. Together, these comparisons suggest that different aspects of these evolving EEG representations of decision variables are related to the neural processes for perception and confidence.

### Neural processes for confidence

The analysis above shows that the EEG representation of accumulated evidence reflected greater differences from the optimal presented evidence *L* in trials where behaviour does not match the optimal response. This suggests that the corresponding neural signals reflect more closely *L\** (the internal evidence actually used by observers to decide) than *L*. To isolate the neural signals which reflect *L*\*, we assume that *L\** approximates *L* with normally distributed errors, and that these errors have larger variance on trials leading to responses that do not match the optimal evidence *L* (a similar approach as in Van Bergen et al., 2015). We used multivariate Bayesian scan statistics (Neill, 2011; Neill, 2019) to cluster signals in space (electrode location) and time where the variance from *L* in the neural representation corresponded to deviations in *L\**, based on behaviour. The statistic tested whether the variability in the neural representation was related to *L\** to a greater extent than could be explained by measurement noise alone (see **Supplementary Note 7** for further details). In this way, the statistic isolates signals more closely related to *L\** than can be explained by *L*, taking into account the noise affecting our measurement of these neural signals.

For perceptual decision-making, signals related to *L\** were initially clustered over posterior electrodes, becoming dispersed over more anterior electrodes late in the epoch (**Figure 5a**, top). For confidence, we found two co-temporal clusters in posterior and anterior electrodes emerging from 668 ms to 824 ms from stimulus onset (**Figure 5a**, bottom). In **Figure 5a** we highlight an early posterior cluster of signals strongly related to *L\** for perceptual decisions, that was not diagnostic of confidence evaluations (in fact the evidence was in favour of the null hypothesis; summed log likelihood ratio = −1176). We obtained cluster-wide representations of *L* from the signals in this early posterior cluster and the two confidence related clusters. The precision of these representations is shown in **Figure 5b**, left. That the information from these clusters is not redundant is evident from the fact that combining the clusters improves the representation precision (**Figure 5b**). For simplicity, we combined the two confidence clusters for further analysis. Similar to the previous analysis (**Figure 4d**), the representation precision of the early posterior cluster was attenuated for the last four samples of the More condition. But, the representation precision of the confidence cluster was maintained (a repeated measures ANOVA revealed a significant interaction between cluster and condition for decoding precision in the last four samples, *F*(1,19) = 32.00, *p* = 0.001, Bonferroni corrected for three comparisons). These results are consistent with dissociable stages of neural processing for confidence evaluation and perceptual decision-making, and support the computational modelling in suggesting a partial dissociation between the internal evidence used for making perceptual decisions and confidence evaluations.

**Figure 5.**
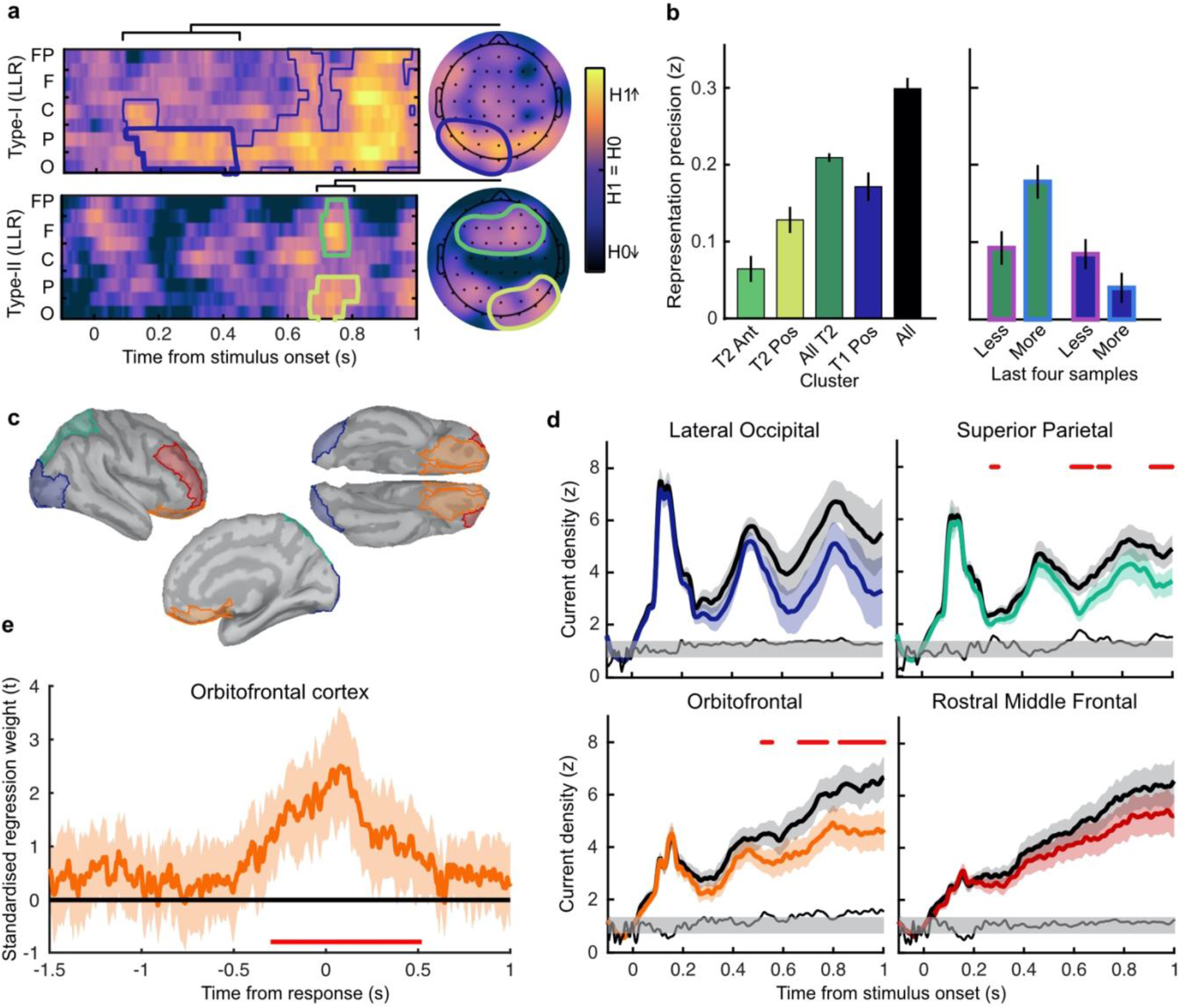
Clusters of behaviourally relevant representations and their sources. **a)** Log likelihood ratio (LLR) of the data given the hypothesis that decoding precision varies with behavioural suboptimalities, against the null hypothesis that decoding precision varies only with measurement noise. Perceptual (Type-I) behaviour is shown on top and confidence (Type-II) behaviour is shown on the bottom. Clusters where the log posterior odds ratio outweighed the prior are circled, only the bold area of the perceptual cluster was further analysed. Time series (left) show the maximum LLR of electrodes laterally, with frontal polar electrodes at the top descending to occipital electrodes at the bottom. Scalp maps (right) show the summed LLR over the indicated time windows. **b)** Left: representation precision (z) training and testing on signals within the clusters. Colours correspond to the circles in a), with the dark green bar showing the combined decoding precision of the anterior and posterior confidence clusters, and the black bar showing the combined representation precision of all clusters. Right: Representation precision of the last four samples in the Less and the More conditions for the combined confidence representation and the perceptual representation. Error bars show 95% within-subject confidence intervals. **c)** ROIs (defined by mindBoggle coordinates; Klein et al., 2017): lateral occipital cortex (blue); superior parietal cortex (green); orbitofrontal cortex (orange); and rostral middle frontal cortex (red). **d)** ROI time series for Noise Max (black) and Noise Min (coloured) epochs, taking the average rectified normalised current density (z) across participants. Shaded regions show 95% within-subject confidence intervals, red horizontal lines indicate cluster corrected significant differences. Standardised within-subject differences are traced above the x-axis, with the shaded region marking z = 0 to z = 1.96 (95% confidence).**e)** Standardised regression weight (t-statistic) of the GLM comparing observers’ confidence ratings to those predicted from the activity localised to the orbitofrontal cortex. The shaded region shows the 95% between subject confidence interval, and the red horizontal line marks the time-window showing cluster-corrected significant differences from 0.

We used the representation from the confidence cluster as an estimate of the internal evidence on which observers base their confidence ratings. We then took the difference from *L* in the estimate of *L\** from the cluster representation as an estimate of the single-sample inference error. This estimate of the single-sample inference error was significantly correlated with the single-sample inference error estimated from the computational model of confidence ratings (*t*(19) = 5.12, *p* < 0.001), and this correlation was significantly greater than the correlation with the error estimated from the model of perceptual decisions alone (*t*(19) = 2.62, *p* = 0.017; see **Supplementary Note 8**). This suggests that this cluster representation is indeed reflecting activity specific to the computation of confidence.

We asked what processes were responsible for driving variability in the internal evidence for confidence beyond what could be explained by the evidence presented to the observer. We selected ‘Noise Min’ and ‘Noise Max’ epochs as the top and bottom quartile of epochs sorted by the estimate of the inference error from the cluster representation, and examined the source-localised EEG activity across these epochs. The presented sensory evidence was similar across Noise Min and Noise Max epochs (see **Supplementary Note 8**), but the additional variability in the Noise Max epochs pushes the represented evidence further from the mean, and should therefore correspond to a greater absolute normalised signal. We estimated the sources of activity in the Noise Min and Noise Max epochs using a template brain (see **Methods**) and tested for differences in the rectified normalised current density in ROIs defined based on the previous literature (**Figure 5c**; Graziano, Parra, and Sigman, 2015; Gherman and Philiastides, 2018; Herding et al., 2019, see **Supplementary Note 9**). As expected, Noise Max epochs showed a greater increase in current density power over time. Significant differences first emerged in the superior parietal cortex (**Figure 5d**; 276 - 304 ms; *t_avg_*(19) = 2.37, *p_cluster_* = 0.016, re-emerging at 596 – 748 ms; *t_avg_*(19) = 2.53, *p_cluster_* = 0.016; and 912 ms; *t_avg_*(19) = 2.50, *p_cluster_* = 0.014), and then in the orbitofrontal cortex (OFC; 516 – 556 ms; *t_avg_*(19) = 2.30, *p_cluster_* = 0.022, re-emerging at 660 – 772 ms; *t_avg_*(19) = 2.79, *p_cluster_* = 0.032, and 824 – 1000 ms; *t_avg_*(19) = 2.60, *p_cluster_* = 0.022). No differences in the rostral middle frontal cortex nor lateral occipital cortex survived cluster correction.

Whilst the activity localised to the superior parietal cortex reflected stimulus driven computations (the consecutive peaks correspond temporally to the response to subsequent stimuli), the activity localised to the orbitofrontal cortex was more indicative of an accumulation process across samples (a smoother increase in signal over time). As an exploratory analysis, we tested whether the activity localised to the orbitofrontal cortex could predict observers’ confidence ratings, presumably by accumulating evidence for evaluating confidence up to the observers’ perceptual decision response. Indeed, the activity localised to the orbitofrontal cortex predicted observers’ confidence ratings, based on the predictions of a generalised linear model with 90/10 cross validation: the standardised regression coefficients increased up to and continued after the perceptual decision response (**Figure 5e**, a significant cluster was located from −300 to 520 ms around the time of the response; *t_ave_*(19) = 3.46, cluster-corrected *p* < 0.001).

## Discussion

We examined the dynamic neural signals associated with the accumulation of evidence for evaluating confidence in perceptual decisions. Observers were required to integrate evidence over multiple samples provided by a sequence of visual stimuli. When observers were unable to control the amount of evidence they were exposed to, they employed a covert decision bound, committing to perceptual decisions when they had enough evidence, even if stimulus presentation continued. We had previously shown evidence for this premature decision commitment based on behaviour and computational modelling (Balsdon, Wyart and Mamassian, 2020). We replicated these results here, and further examined the neural signatures of covert decision making. We found that the distribution of spectral power associated with the preparation and execution of motor responses in the Free task (where the response is entered as soon as the decision is made) could be used to accurately predict responses in the More condition of the Replay task over 1 s prior to when the response was entered, and with significantly greater sensitivity than in the Less condition (when observers were unlikely to have committed to a decision early). This suggests that covert decisions could trigger the motor preparation for pressing the response key. Moreover, the strength of the eventual motor response signal could be predicted by earlier decision evidence in the More condition, as if observers are maintaining some representation of the decision evidence whilst waiting to press the response key.

Based on the evoked representation of accumulated evidence, perceptual decision accuracy relied on a flow of information processing from early occipital and parietal signals, which then spread through to anterior electrodes. When observers committed to perceptual decisions prematurely, only the early part of the representation of accumulated evidence was attenuated. This selective dampening of the representation of accumulated evidence following premature decision commitment delineates which computations are devoted solely to the perceptual decision process, and which computations reflect the input to the decision process: The representations of stimulus orientation and decision update (Wyart et al., 2012; Wyart et al., 2015; Weiss et al., 2021), which are necessary input for the perceptual decision, did not substantially change after committing to a perceptual decision. This initial perceptual processing stage likely remained important for the continued accumulation of evidence for evaluating confidence (even after the completion of perceptual decision processes), though it could also be that these processes are automatically triggered by stimulus onset irrespective of whether the evidence is being accumulated for decision-making.

Confidence should increase with increasing evidence for the perceptual decision. It is therefore unsurprising that the neural correlates of confidence magnitude have found similar EEG markers as those related to the accumulation of the underlying perceptual decision evidence: the P300 (Gherman and Philiastides, 2015; Desender et al., 2016; Desender et al., 2019; Zakrzewski et al., 2019; Rausch et al., 2020); and Central Parietal Positivity (CPP; Boldt et al., 2019; Herding et al., 2019, indeed we show a similar effect in **Supplementary Note 4**). The analysis presented in this manuscript targeted confidence precision rather than confidence magnitude, by assessing confidence relative to an optimal observer who gives high confidence ratings on trials where the evidence in favour of the perceptual choice is greater than the median across trials. We isolated part of the neural representation of accumulated evidence where imprecision relative to the optimal presented evidence predicted greater deviations from optimal in the internal representation of evidence used for confidence evaluation implied from behaviour. The internal evidence predicted from this neural representation was also more strongly related to the evidence for confidence than the evidence used for perceptual decisions based on the computational model fit to describe behaviour.

We analysed the sources of activity more closely representing the internal evidence on which the confidence evaluation was based than the optimal presented evidence. Activity localised to the Superior Parietal and Orbitofrontal cortices was found to track this internal evidence for confidence throughout decision-making. This is not at odds with the previous literature: The difference in superior parietal cortex could be linked with findings from electrophysiology that suggest that confidence is based on information coded in parietal cortex, where the underlying perceptual decision evidence is integrated (Kiani et al., 2009; Rutishauser et al., 2018; though at least a subset of these neurons reflect bounded accumulation, which is in contrast with the continued confidence accumulation described in this experiment; Kiani, Hanks, and Shadlen, 2007). Early electrophysiological investigation into the function of the orbitofrontal cortex revealed neural coding associated with stimulus value (Thorpe, Rolls, and Maddison, 1983), which has since been linked with a confidence-modulated signal of outcome-expectation (Kepecs et al., 2008; and in human fMRI; Rolls, Grabenhorst, and Deco, 2010) and recently, shown to be domain-general (single OFC neurons were associated with confidence in both olfactory and auditory tasks; Masset et al., 2020). The source localisation analysis therefore connects previous findings, indicating confidence feeds off an evidence accumulation process, culminating in higher-order brain areas that appear to function for guiding outcome-driven behaviour based on decision certainty.

These neural signatures of confidence evidence encoding were present throughout the process of making a perceptual decision. This is in line with more recent evidence suggesting that confidence could be computed online, alongside perceptual evidence accumulation (Zizlsperger et al., 2014; Gherman and Philiastides, 2015; Balsdon et al., 2020), as opposed to assessing the evidence in favour of the perceptual decision only after committing to that decision. Computational model comparison supported this interpretation, showing the best description of confidence behaviour was an accumulation process that was partially dissociable from perceptual evidence accumulation (**Supplementary Note 1**; replicating our previous analysis, Balsdon et al., 2020). This partial dissociation mediates the ongoing debate between single-channel (for example, Maniscalco and Lau, 2016) and dual-channel (for example, Charles, King, and Deheane 2014) models, as it constrains confidence by perceptual suboptimalities, at the same time as allowing additional processing to independently shape confidence. The combination of this partial dissociation and online monitoring could allow for metacognitive control of perceptual evidence accumulation, to flexibly balance perceptual accuracy against temporal efficiency, by bounding perceptual evidence accumulation according to contemporaneous confidence.

Using this protocol, we were able to delineate two distinct representations of accumulated evidence which correspond to perceptual decision-making and confidence evaluations. These neural representations were partially dissociable in that the perceptual representation neglected additional evidence following premature decision commitment whilst the confidence representation continued to track the updated evidence independently of decision commitment. This partial dissociation validates the predictions of the computational model and provides a framework for the cognitive architecture underlying the distinction between perception and confidence. That the neural resources involved in the confidence representation can be recruited independently of perceptual processes implies a specific neural circuit for the computation of confidence, a necessary feature of a general metacognitive mechanism flexibly employed to monitor the validity of any cognitive process.

## Methods

### Participants

A total of 20 participants were recruited from the local cognitive science mailing list (RISC) and by word of mouth. No participant met the pre-registered (https://osf.io/346pe/?view_only=ddbc092996f34438964cf45a239498bb) exclusion criteria of chance-level performance or excessive EEG noise. Written informed consent was provided prior to commencing the experiment. Participants were required to have normal or corrected to normal vision. Ethical approval was granted by the INSERM ethics committee (ID RCB: 2017-A01778-45 Protocol C15-98).

### Materials

Stimuli were presented on a 24” BenQ LCD monitor running at 60 Hz with resolution 1920×1080 pixels and mean luminance 45 cd/m^2^. Stimulus generation and presentation was controlled by MATLAB (Mathworks) and the Psychophysics toolbox (Brainard, 1997; Pelli, 1997; Kleiner et al., 2007), run on a Dell Precision M4800 Laptop. Observers viewed the monitor from a distance of 57 cm, with their head supported by a chin rest. EEG data were collected using a 64-electrode BioSemi ActiveTwo system, run on a dedicated mac laptop (Apple Inc.), with a sample rate of 512 Hz. Data were recorded within a shielded room.

### Stimuli

Stimuli were oriented Gabor patches displayed at 70% contrast, subtending 4 dva and with spatial frequency 2 cyc/deg. On each trial a sequence of stimuli was presented, at an average rate of 3 Hz, with the stimulus presented at full 70% contrast for a variable duration between 50 and 83 ms, with a sudden onset, followed by an offset ramp over two flips, where the stimulus contrast decreased by 50% and 75% before complete offset. Stimulus onset timing was jittered within the stimulus presentation interval such that the timing of stimulus onset was irregular but with at least 216 ms between stimuli. These timings and stimulus examples are shown in **Figure 1a**.

On each trial the orientations of the presented Gabors were drawn from one of two circular Gaussian (Von Mises) distributions centred on +/− 45° from vertical (henceforth referred to as the ‘orange’ and ‘blue’ distributions respectively), with concentration κ = 0.5 (shown in **Figure 1d**). Stimuli were displayed within an annular ‘colour-guide’ where the colour of the annulus corresponds to the probability of the orientation under each distribution, using the red and blue RGB channels to represent the probabilities of each orientation under each distribution. Stimuli were presented in the centre of the screen, with a black central fixation point to guide observers’ gaze.

### Procedure

The task was a modified version of the weather prediction task (Knowlton et al., 1996; Drugowitsch et al., 2016). Throughout the experiment, the observer’s perceptual task was to categorise which distribution the stimulus orientations were sampled from. They were instructed to press the ‘d’ key with their left hand (of a standard querty keyboard) for the blue distribution and the ‘k’ key with their right hand for the orange distribution. There were two variants of the task: The Free task and the Replay task. The trials were composed of three repetitions of 100 predefined sequences of up to 40 samples (50 trials from each distribution) for each observer (300 trials per task).

In the ‘Free’ task, observers were continually shown samples (up to 40) until they entered their response. They were instructed to enter their response as soon as they ‘feel ready’ to make a decision, with emphasis on both accuracy (they should make their decision when they feel they have a good chance of being correct) and on time (they shouldn’t take too long to complete each trial). A graphical description of this task is shown in **Figure 1b**.

After completing the Free task, observers then completed the Replay task. In this task they were shown a specific number of samples and could only enter their response after the sequence finished, signalled by the fixation point turning red. The number of samples was determined based on the number observers chose to respond to in the Free task. There were three intermixed conditions: In the Less condition observers were shown two fewer samples than the minimum they had chosen to respond to on that predefined sequence in the Free task; In the Same condition observers were shown the median number of samples from that predefined sequence; in the More condition observers were shown four additional samples compared to the maximum number they chose to respond to on that sequence in the Free task. After entering their perceptual (Type-I) response, observers were cued to give a confidence rating (Type-II decision). The confidence rating was given on a 4-point scale where 1 represents very low confidence that the perceptual decision was correct, and 4, certainty that the perceptual decision was correct. The rating was entered by pressing the ‘space bar’ when a presented dial reached the desired rating. The dial was composed of a black line which was rotated clockwise to each of 4 equidistant angles (marked 1 - 4) around a half circle, at a rate of 1.33 Hz. The dial started at a random confidence level on each trial and continued updating until a rating was chosen. A graphical description of this task is shown in **Figure 1c**.

Prior to commencing the experimental trials, participants were given the opportunity to practice the experiment and ask questions. They first performed 20 trials of a fixed number of samples with only the perceptual decision, with feedback after each response as to the true category. They then practiced the Replay task with the confidence rating (and an arbitrary number of samples). Finally, they practiced the Free task, before commencing the experiment with the Free task.

## Analysis

### Behaviour

Perceptual (Type-I) decisions were evaluated relative to the category the orientations were actually drawn from. Performance is presented as proportion correct, whilst statistical analyses were performed on sensitivity (d’). Sensitivity was calculated based on the proportion of hits (responding “Category A” when category A was presented) and false alarms (responding “Category A” when category B was presented). Confidence was evaluated relative to an optimal observer who gives high confidence when the log-likelihood of the chosen category, based on the presented orientations, is above the median across trials, and low confidence on trials with less than the median log-likelihood. More broadly, confidence should increase with increasing evidence in favour of the perceptual decision, see **Supplementary Note 3**. A General Linear Model was used to validate the influence of the optimal presented evidence on perceptual decisions and confidence evaluations. The accumulated evidence up to the final sample and four samples before the response was used as a regressor for the perceptual decision assuming a binomial distribution with a probit link function. A comparable analysis was performed for confidence by binarizing confidence ratings into Low (ratings of 1 or 2) and High (ratings of 3 or 4) and taking the evidence signed by the perceptual decision.

### Computational modelling

Computational modelling followed the same procedure as Balsdon, Wyart, and Mamassian (2020). The model parametrically describes suboptimalities relative to the Bayesian optimal observer. The Bayesian optimal observer knows the category means, 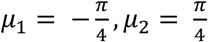, and the concentration, κ = 0.5, and takes the probability of the orientation *θ*_*n*_ (at sample *n*) given each category *ψ* (*ψ* = 1 or *ψ* = 2)

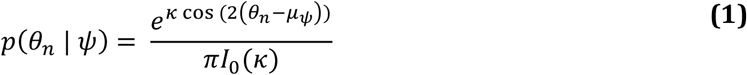

where *I*_0_(.) is the modified Bessel function of order 0. The optimal observer then chooses the category *ψ* with the greatest posterior probability over all samples for that trial, *T* (*T* varies from trial to trial). Given a uniform category prior, 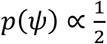, and perfect anticorrelation in *p*(*θ*_*n*_ │ *ψ*) over the categories, the log posterior is proportional to the sum of the difference in the log-likelihood for each category (*ℓ*_*n*_ = *ℓ*_*n*,1_ – *ℓ*_*n*,2_)

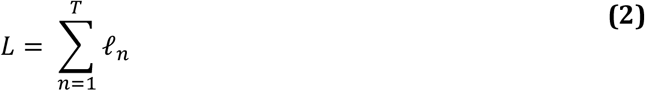

where:

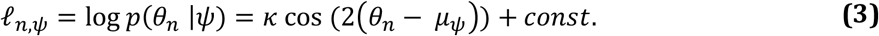

Such that the Bayesian optimal decision is 1 if *z* > 0 and 2 if *z* ≤ 0.

The suboptimal observer suffers inaccuracies in the representation of each evidence sample, captured by additive independent identically distributed (i.i.d) noise, *ε*_*n*_. The noise is Gaussian distributed with zero mean, and the degree of variability parameterised by σ, the standard deviation

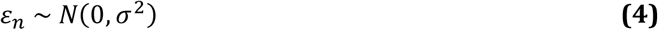

The evidence over samples is also imperfectly accumulated, incurring primacy or recency biases parameterised by *α*, the weight on the current accumulated evidence compared to the new sample (*α* > 1 creates a primacy effect). By the end of the trial, the weight on each sample *n* is equal to

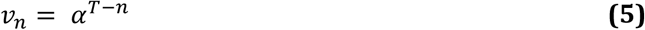

where *T* is the eventual total samples on that trial and *n* ∈ [1, *T*].

In the Free task the observer responds when accumulated evidence reaches a bound, Λ. The optimal observer sets a constant bound on proportion correct over sequence length, which is an exponential function on the average evidence over the samples accumulated. The human observer can set the scale, *b*, and the rate of decline, *λ*, of the bound suboptimally, resulting in

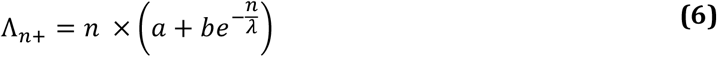

for the positive decision bound (the negative bound, Λ_*n*−_ = –Λ_*n*+_). The likelihood *f*(*n*) of responding at sample n was estimated by computing the frequencies, over 1000 samples from *ε*_*n*_ (Monte Carlo simulation), of first times where the following inequality is verified

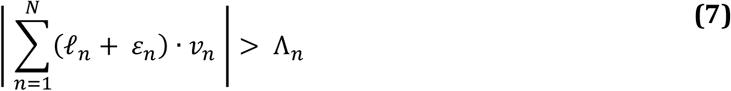

The response time, relative to reaching the decision bound, is delayed by non-decision time for executing the motor response, which is described by a Gaussian distribution of mean, *μ*_*U*_, and variance, σ_*U*_^2^.

### Model fitting

Parameters were optimised to minimise the negative log-likelihood of the observer making response *r* on sample *n* on each trial for each participant using Bayesian Adaptive Direct Search (Acerbi and Ma, 2017). The log-likelihoods were estimated using Monte Carlo Simulation, with the sensitivity of this approach being addressed in previous work (Balsdon et al., 2020). The full model was simplified using a knock-out procedure based on Bayesian Model Selection (Rigoux et al., 2014) to fix the bias (exceedance probability = 0.93) and lapse (exceedance probability >0.99) parameters (not described above, see **Supplementary Note 1**).

In the Replay task, confidence ratings were fit using the same model described above, but with additional criteria determining confidence ratings, described by three bounds on the confidence evidence, parameterised in the same manner as the decision bound. These models were then used to simulate the internal evidence of each observer from sample to sample, and the error compared to the optimal evidence (uncorrupted by suboptimalities, see **Supplementary Note 2**).

### EEG pre-processing

EEG data were pre-processed using the PREP processing pipeline (Bigdely-Shamlo, et al., 2015), implemented in EEGlab (v2019.0, Delorme & Makeig, 2004) in MATLAB (R2019a, Mathworks). This includes line noise removal (notch filter at 50 Hz and harmonics) and re-referencing (robust average re-reference on data detrended at 1 Hz). The data were then filtered to frequencies between 0.5 and 80 Hz, and down-sampled to 256 Hz. Large epochs were taken locked to each stimulus (−500 to 1500 ms) and each response (−5000 to 1500 ms). Independent Components Analysis was used to remove artefacts caused by blinks and excessive muscle movement identified using labels with a probability greater than 0.35 from the ICLabel project classifier (Swartz Centre for Computational Neuroscience).

### Response classification analysis

The power spectrum across frequency tapers from 1 to 64 Hz with 25% spectral smoothing was resolved using wavelet convolution implemented in FieldTrip (Oostenveld et al., 2011). The epochs were then clipped at −3 to 1 s around the time of entering the perceptual response. Linear discriminant analysis was performed to classify perceptual responses, using 10-fold cross validation, separately on each taper at each time-point. An analysis of the frequencies contributing to accurate classification at the time of the response revealed significant contributions from 8 to 26 Hz (**Supplementary Note 4**). We therefore continued by using the power averaged across these frequency bands to train and test the classifier. Classifier accuracy was assessed using the area under the receiver operating characteristic curve (AUC). At the single-trial level, the probability of the response based on the classifier was computed from the relative normalised Euclidean distance of the trial features from the response category means in classifier decision space.

### Encoding Variable Regression

We used a linear regression analysis to examine the EEG correlates of different aspects of the decision evidence (encoding variables) in epochs locked to stimulus onset. Regularised ridge regression (ridge λ = 1) was used to predict the encoding variables based on EEG data, over 10-fold cross validation. The precision of the representation of each encoding variable was computed within each observer by taking the Fisher transform of the correlation coefficient (Pearson’s r) between the encoded variable and predicted variable. To maximise representation precision, the data were bandpass filtered (1 – 8 Hz) and decomposed into real and imaginary parts using a Hilbert Transform (**Supplementary Note 5**). For each time point, the data from all electrodes were used to predict the encoded variable. The temporal generalisation of decoding weights was examined by training at one time point and testing at another. The contribution of information from signals at each electrode was examined by training and testing on the signals at each electrode at each time point (further details in **Supplementary Note 5**).

Behaviourally relevant signals were isolated by comparing representation precision at each time point and electrode for epochs leading to optimal perceptual and confidence responses, compared to responses that did not match the optimal observer. Cluster modelling was used to isolate contiguous signals where the log posterior odds were in favour of the alternative hypothesis that the representation systematically deviated further from the optimal presented evidence than what could be explained by measurement noise alone (**Supplementary Note 6**). New regression weights were then calculated on signals from the entire cluster and representation errors calculated as the difference of the predicted variable from the expected value given the representation.

### Source Localisation

Identifying the clusters of signals associated with confidence processes offers relatively poor spatial and temporal (given the bandpass filter; de Cheveigné, and Nelken, 2019) resolution for identifying the source of confidence computations. Source localisation was therefore performed, using Brainstorm (Tadel et al., 2011). The forward model was computed using OpenMEEG (Gramfort et al., 2010; Kybic et al., 2005) and the ICBM152 anatomy (Fonov et al., 2011; 2009). Two conditions were compared, Noise Min and Noise Max, which corresponded to quartiles of epochs sorted by representation error in the confidence clusters (see **Supplementary Note 7** for more details). Cortical current source density was estimated from the average epochs using orientation-constrained minimum norm imaging (Baillet, Mosher, and Leahy, 2001). ROIs in the Lateral Occipital, Superior Parietal, Rostral Middle Frontal (including dlPFC), Medial Orbitofrontal, and rostral Anterior Cingulate Cortex, were defined using MindBoggle coordinates (Klein et al., 2017). Statistical comparisons were performed on the bilateral ROI time series (using cluster correction and a minimum duration of 20 ms), computed over separate conditions on rectified normalised subject averages (low-pass filtered at 40 Hz).

To predict confidence magnitude from the activity localised to the orbitofrontal cortex, we recovered to current density from 20 subregions (approximately equal parcellations) of the orbitofrontal cortex in epochs locked to the time of the response. A general linear model (assuming a normal distribution with identity link) was used to predict the observers’ confidence ratings on held-out data (90/10 cross-fold) from the neural activity at each time-point leading to the response. The prediction was quantified as the standardised regression weight from a new general linear model comparing the predicted and actual confidence ratings across all folds.

## References

Acerbi, L., & Ma, W. J. Practical Bayesian optimization for model fitting with Bayesian adaptive direct search. In Advances in Neural Information Processing Systems, December 2017; 1836–1846

Bahrami, B., Olsen, K., Bang, D., Roepstorff, A., Rees, G., & Frith, C. What failure in collective decision-making tells us about metacognition. Philosophical Transactions of the Royal Society B: Biological Sciences, 2012; 367(1594), 1350–1365.

Baillet, S., Mosher, J. C., & Leahy, R. M. Electromagnetic brain mapping. IEEE Signal Processing Magazine, 2001; 18(6), 14–30.

Balsdon, T., Wyart, V., & Mamassian, P. Confidence controls perceptual evidence accumulation. Nature Communications, 2020; 11(1), 1–11

Bang, J. W., Shekhar, M., & Rahnev, D. Sensory noise increases metacognitive efficiency. Journal of Experimental Psychology: General, 2019; 148(3), 437.

Baranski, J. V., & Petrusic, W. M. The calibration and resolution of confidence in perceptual judgments. Perception & Psychophysics, 1994; 55(4), 412–428.

Bigdely-Shamlo, N., Mullen, T., Kothe, C., Su, K. M., & Robbins, K. A. The PREP pipeline: standardized preprocessing for large-scale EEG analysis. Frontiers in Neuroinformatics, 2015; 9, 16

Boldt, A., Schiffer, A. M., Waszak, F., & Yeung, N. Confidence predictions affect performance confidence and neural preparation in perceptual decision making. Scientific Reports, 2019; 9(1), 1–17.

Ruby, E., Maniscalco, B., & Peters, M. A. On a ‘failed’ attempt to manipulate visual metacognition with transcranial magnetic stimulation to prefrontal cortex. Consciousness and cognition, 2018; 62, 34–41.

Brainard, D. H. The psychophysics toolbox. Spatial Vision, 1997; 10(4), 433–436.

Charles, L., King, J. R., & Dehaene, S. Decoding the dynamics of action, intention, and error detection for conscious and subliminal stimuli. Journal of Neuroscience, 2014; 34(4), 1158–1170.

Cortese, A., Amano, K., Koizumi, A., Kawato, M., & Lau, H. Multivoxel neurofeedback selectively modulates confidence without changing perceptual performance. Nature communications, 2016; 7(1), 1–18.

de Cheveigné, A., & Nelken, I. Filters: when, why, and how (not) to use them. Neuron, 2019; 102(2), 280–293.

Delorme, A., & Makeig, S. EEGLAB: an open source toolbox for analysis of single-trial EEG dynamics including independent component analysis. Journal of Neuroscience Methods, 2004; 134(1), 9–21.

Denison, R. N., Adler, W. T., Carrasco, M., & Ma, W. J. Humans incorporate attention-dependent uncertainty into perceptual decisions and confidence. Proceedings of the National Academy of Sciences, 2018 115(43), 11090–11095.

Desender, K., Van Opstal, F., Hughes, G., & Van den Bussche, E. The temporal dynamics of metacognition: Dissociating task-related activity from later metacognitive processes. Neuropsychologia, 2016; 82, 54–64.

Desender, K., Murphy, P., Boldt, A., Verguts, T., & Yeung, N. A postdecisional neural marker of confidence predicts Information-Seeking in Decision-Making. Journal of Neuroscience, 2019; 39(17), 3309–3319.

Drugowitsch, J., Wyart, V., Devauchelle, A. D., & Koechlin, E. Computational precision of mental inference as critical source of human choice suboptimality. Neuron, 2016; 926, 1398–1411

Fleming, S. M., Huijgen, J., & Dolan, R. J. Prefrontal contributions to metacognition in perceptual decision making. Journal of Neuroscience, 2012; 32(18), 6117–6125.

Fleming, S. M., Ryu, J., Golfinos, J. G., & Blackmon, K. E. Domain-specific impairment in metacognitive accuracy following anterior prefrontal lesions. Brain, 2014; 137(10), 2811–2822.

Fleming, S. M., & Daw, N. D. Self-evaluation of decision-making: A general Bayesian framework for metacognitive computation. Psychological Review, 2017; 124(1), 91.

Fleming, S. M., Van Der Putten, E. J., & Daw, N. D. Neural mediators of changes of mind about perceptual decisions. Nature neuroscience, 2018; 21(4), 617–624.

Fonov VS, Evans AC, McKinstry RC, Almli CR, Collins DL. Unbiased nonlinear average age-appropriate brain templates from birth to adulthood. NeuroImage, 2009; 47, S102.

Fonov, V., Evans, A. C., Botteron, K., Almli, C. R., McKinstry, R. C., Collins, D. L., & Brain Development Cooperative Group. Unbiased average age-appropriate atlases for pediatric studies. Neuroimage, 2011; 54(1), 313–327.

Frith, C. D. The role of metacognition in human social interactions. Philosophical Transactions of the Royal Society B: Biological Sciences, 2012; 367(1599), 2213–2223.

Gherman, S., & Philiastides, M. G. Human VMPFC encodes early signatures of confidence in perceptual decisions. eLife, 2018; 7, e38293.

Gherman, S., & Philiastides, M. G. Neural representations of confidence emerge from the process of decision formation during perceptual choices. NeuroImage, 2015; 106, 134–143.

Gramfort, A., Papadopoulo, T., Olivi, E., & Clerc, M. OpenMEEG: opensource software for quasistatic bioelectromagnetics. Biomedical Engineering Online, 2010; 9(1), 45.

Graziano, M., Parra, L. C., & Sigman, M. Neural correlates of perceived confidence in a partial report paradigm. Journal of Cognitive Neuroscience, 2015; 27(6), 1090–1103.

Geurts, L. S., Cooke, J. R., van Bergen, R. S., & Jehee, J. F. Subjective confidence reflects representation of Bayesian probability in cortex. 2021; bioRxiv.

Helmholtz, H.L.F.v. Treatise on Physiological Optics, Thoemmes Press 1856.

Herding, J., Ludwig, S., von Lautz, A., Spitzer, B., & Blankenburg, F. Centro-parietal EEG potentials index subjective evidence and confidence during perceptual decision making. NeuroImage, 2019; 201, 116011.

Kepecs, A., Uchida, N., Zariwala, H. A., & Mainen, Z. F. Neural correlates, computation and behavioural impact of decision confidence. Nature, 2008; 455, 227–231.

Kiani, R., & Shadlen, M. N. Representation of confidence associated with a decision by neurons in the parietal 772 cortex. Science, 2009; 324, 759–764.

Kiani, R., Corthell, L., & Shadlen, M. N. Choice certainty is informed by both evidence and decision time. Neuron, 2014; 84(6), 1329–1342.

Kiani, R., Hanks, T. D., & Shadlen, M. N. Bounded integration in parietal cortex underlies decisions even when viewing duration is dictated by the environment. Journal of Neuroscience, 2008; 28(12), 3017–3029.

Klein, A., Ghosh, S. S., Bao, F. S., Giard, J., Häme, Y., Stavsky, E., … &Keshavan, A. Mindboggling morphometry of human brains. PLoS Computational Biology, 2017; 13(2), e1005350.

Kleiner, M., Brainard, D., & Pelli, D. What’s new in Psychtoolbox-3? 2007.

Knowlton, B. J., Mangels, J. A., & Squire, L. R. A neostriatal habit learning system in humans. Science, 1996; 273(5280), 1399–1402

Kybic, J., Clerc, M., Abboud, T., Faugeras, O., Keriven, R., & Papadopoulo, T. A common formalism for the integral formulations of the forward EEG problem. IEEE Transactions on Medical Imaging, 2005; 24(1), 12–28.

Lak, A., Costa, G. M., Romberg, E., Koulakov, A. A., Mainen, Z. F., & Kepecs, A. Orbitofrontal cortex is required for optimal waiting based on decision confidence. Neuron, 2014; 84(1), 190–201.

Lapate, R. C., Samaha, J., Rokers, B., Postle, B. R., & Davidson, R. J. Perceptual metacognition of human faces is causally supported by function of the lateral prefrontal cortex. Communications biology, 2020; 3(1), 1–10.

Maniscalco, B., & Lau, H. The signal processing architecture underlying subjective reports of sensory awareness. Neuroscience of Consciousness, 2016; 1.

Masset, P., Ott, T., Lak, A., Hirokawa, J., & Kepecs, A. Behavior- and modality-general representation of confidence in orbitofrontal cortex. Cell, 2020; 182(1), 112–126.

Mazancieux, A., Fleming, S., Souchay, C., & Moulin, C. Retrospective confidence judgments across tasks: domain-general processes underlying metacognitive accuracy. BioRxiv 2018.

Moreno-Bote, R. Decision confidence and uncertainty in diffusion models with partially correlated neuronal 797 integrators. Neural Computation, 2010; 22, 1786–1811.

Murphy, P. R., Robertson, I. H., Harty, S., & O’Connell, R. G. Neural evidence accumulation persists after choice to inform metacognitive judgments. Elife, 2015; 4, e11946.

Neill, D. B. Fast Bayesian scan statistics for multivariate event detection and visualization. Statistics in Medicine, 2011; 30(5), 455–469.

Neill, D. B. Bayesian Scan Statistics. In: Glaz J., Koutras M. (eds) Handbook of Scan Statistics. Springer, New York, NY. 2019.

Oostenveld, R., Fries, P., Maris, E., & Schoffelen, J. M. FieldTrip: open source software for advanced analysis of MEG, EEG, and invasive electrophysiological data. Computational Intelligence and Neuroscience, 2011.

Pelli, D. G. The VideoToolbox software for visual psychophysics: Transforming numbers into movies. Spatial Vision, 1997; 10, 437–442.

Pleskac, T. J., & Busemeyer, J. R. Two-stage dynamic signal detection: a theory of choice, decision time, and confidence. Psychological Review, 2010; 117(3), 864.

Pollack, I., & Decker, L. R. Confidence ratings, message reception, and the receiver operating characteristic. The Journal of the Acoustical Society of America, 1958; 30(4), 286–292.

Ratcliff, R. A theory of memory retrieval. Psychological Review, 1987; 85(2), 59.

Rausch, M., Zehetleitner, M., Steinhauser, M., & Maier, M. E. Cognitive modelling reveals distinct electrophysiological markers of decision confidence and error monitoring. NeuroImage, 2020; 218, 116963.

Rigoux, L., Stephan, K.E., Friston, K.J. & Daunizeau, J. Bayesian Model Selection for Group Studies Revisited. NeuroImage 2014; 84, 971–85.

Rolls, E. T., Grabenhorst, F., & Deco, G. Choice, difficulty, and confidence in the brain. NeuroImage, 2010; 53(2), 694–706.

Rounis, E., Maniscalco, B., Rothwell, J. C., Passingham, R. E., & Lau, H. Theta-burst transcranial magnetic stimulation to the prefrontal cortex impairs metacognitive visual awareness. Cognitive neuroscience, 2010; 1(3), 165–175.

Rutishauser, U., Aflalo, T., Rosario, E. R., Pouratian, N., & Andersen, R. A. Single-neuron representation of memory strength and recognition confidence in left human posterior parietal cortex. Neuron, 2018; 97(1), 209–220.

Shekhar, M., & Rahnev, D. Sources of Metacognitive Inefficiency. Trends in Cognitive Sciences.

Tadel, F., Baillet, S., Mosher, J. C., Pantazis, D., & Leahy, R. M. Brainstorm: a user-friendly application for MEG/EEG analysis. Computational Intelligence and Neuroscience, 2011.

Thorpe, S. J., Rolls, E. T., & Maddison, S. The orbitofrontal cortex: neuronal activity in the behaving monkey. Experimental Brain Research, 1983; 49(1), 93–115.

Vaccaro, A. G., & Fleming, S. M. Thinking about thinking: A coordinate-based meta-analysis of neuroimaging studies of metacognitive judgements. Brain and neuroscience advances, 2018; 2, 2398212818810591.

Veenman, M. V., Wilhelm, P., & Beishuizen, J. J. The relation between intellectual and metacognitive skills from a developmental perspective. Learning and Instruction, 2004; 14(1), 89–109.

Vickers, D. Evidence for an accumulator model of psychophysical discrimination. Ergonomics, 1970; 13(1), 37–58.

Vickers, D. Decision processes in visual perception. New York, NY: Academic Press. 1979.

Weiss, A., Chambon, V., Lee, J. K., Drugowitsch, J., & Wyart, V. Interacting with volatile environments stabilizes hidden-state inference and its brain signatures. Nature communications, 2021; 12(1), 1–17.

Wyart, V., De Gardelle, V., Scholl, J., & Summerfield, C. Rhythmic fluctuations in evidence accumulation during decision making in the human brain. Neuron, 2012; 76(4), 847–858.

Wyart, V., Myers, N. E., & Summerfield, C. Neural mechanisms of human perceptual choice under focused and divided attention. Journal of Neuroscience, 2015; 35(8), 3485–3498.

Yokoyama, O., Miura, N., Watanabe, J., Takemoto, A., Uchida, S., Sugiura, M.,&Nakamura, K. Right frontopolar cortex activity correlates with reliability of retrospective rating of confidence in short-term recognition memory performance. Neuroscience research, 2010; 68(3), 199–206.

Zakrzewski, A. C., Wisniewski, M. G., Iyer, N., … & Simpson, B. D. Confidence tracks sensory-and decision-related ERP dynamics during auditory detection. Brain and Cognition, 2019; 129, 49–58

Zizlsperger, L., Sauvigny, T., Händel, B., & Haarmeier, T. Cortical representations of confidence in a visual perceptual decision. Nature Communications, 2014; 5(1), 1–13.

