## Supplemental for "Separable neural signatures of confidence during perceptual decisions"

Supplementary materials

### Supplementary Note 1

##### Computational Model fitting

The computational model is described in full in the **Methods** section. Briefly, the model is based on the Bayesian optimal observer with full knowledge of the category distributions (means $\mu_1$ and $\mu_2$, concentration $\kappa$), and takes as evidence the difference in the log posterior probability ($\ell_n$) of each category given the orientation ($\theta_n$)

$$\ell_n = \ell_{n,1} - \ell_{n,2} = \kappa\cos\big(2(\theta_n - \mu_1)\big) - \kappa\cos\big(2(\theta_n - \mu_2)\big) \qquad \textbf{(1)}$$
$$= 2\,\kappa\sin(\mu_1 - \mu_2)\sin(2\theta_n - \mu_1 - \mu_2) = \sin(2\theta_n),$$

where chosen values ($\kappa = 0.5$, $\mu_1 = -\pi/4$, and $\mu_2 = \pi/4$) have been implemented in the last equation. Whilst the optimal observer perfectly sums the evidence over each sample, the suboptimal human observer accumulates evidence with some temporal integration bias, $\alpha$ (where $\alpha > 1$ creates a primacy effect, and $\alpha < 1$, a recency effect), and incurs inference error (noise in the estimate of the true evidence) parameterised by $\sigma$, the standard deviation of the Gaussian distribution from which each sample of noise, $\varepsilon_n$, is drawn from. The human observer may also experience some response bias, *c* (the tendency to choose one category irrespective of the evidence), and incur lapses (pressing a random key), described by the lapse rate, *l*. The accumulated evidence, $L^*$, up to sample *n*, is suboptimally accumulated by

$$L_n^* = \alpha L_{n-1}^* + \ell_n + \varepsilon_n \qquad \textbf{(2)}$$

The observer then chooses category 1 if $L^* > c$, except on a proportion of trials, *l*, where the response is randomly selected.

These four parameters were used to capture the differences in the human observers' responses (category choice and confidence rating) from the optimal observer who perfectly integrates all evidence presented.

In the Free task, the model was designed not only to describe the category choice, but at which sample the human observer chose to respond. This was achieved via a decision boundary, the nature of which has been addressed in previous work (Balsdon, Wyart, and Mamassian, 2020). The boundaries, $\Lambda_{n+}$ and $\Lambda_{n-}$, follow an exponential function on the average evidence over samples (which is a constant bound on the probability of a correct response), described by three parameters: the minimum, *a*, the scale, *b*, and the rate of decline, $\lambda$

$$\Lambda_{n+} = n \times \left(a + be^{-\frac{n}{\lambda}}\right) \qquad \textbf{(3)}$$

There is an optimal combination of these parameters to achieve any particular proportion correct across the experiment, but the human observer may set their bound suboptimally. In addition, non-decision time (the time from the last sample integrated to pressing the response key) was described by a Normal distribution with mean $\mu_U$, and variance $\sigma_U^2$. Giving an additional five parameters for describing when the observer enters their response.

We followed the same procedure as in Balsdon et al., 2020, involving four stages:

1. Reduce the number of free parameters with a knock-out procedure.
2. Compare (covert) Bound and No-bound models of the perceptual decision in the Replay task.
3. Identify any systematic differences in the parameters required to describe the confidence ratings, compared to the perceptual decision, in order to discern the relationship between processes for perceptual decisions and confidence.
4. Apply the same Bound vs. No-bound comparison for describing the confidence ratings.

The average parameter values and fit metrics for Stage 1. are shown in Table 1. According to this analysis, the bias (c) and lapse rate (*l*) were fixed. There was some evidence the boundary minimum (*a*) could be fixed in the Replay task, but the preference in the Free task was to leave it free to vary.

| **Free Task** | | | | | | | | | | | |
|---|---|---|---|---|---|---|---|---|---|---|---|
| **Model** | $\sigma$ | $\alpha$ | $c$ | $\mu_U$ | $\sigma_U^2$ | $a$ | $b$ | $\lambda$ | $l$ | **LLH** | **ΣBIC** |
| Full | 0.83 | 0.98 | -0.04 | 425 | 0.52 | 0.10 | 6.04 | 1.93 | 0.016 | -734.91 | 30423.01 |
| $\alpha = 1$ | 0.83 | 1.00 | 0.00 | 430 | 0.50 | 0.13 | 6.61 | 2.03 | 0.014 | -734.97 | 30311.59 |
| $c = 0$ | 0.80 | 0.92 | 0.00 | 452 | 0.54 | 0.11 | 5.28 | 2.01 | 0.017 | -736.86 | 30387.02 |
| $\mu_U = 400$ | 0.76 | 0.94 | 0.00 | 400 | 0.52 | 0.09 | 5.52 | 2.23 | 0.016 | -739.77 | 30503.40 |
| $\sigma_U^2 = 1$ | 0.69 | 0.96 | -0.02 | 435 | 1.00 | 0.10 | 6.34 | 1.97 | 0.015 | -754.18 | 31079.84 |
| $a = 0.1$ | 0.77 | 0.92 | 0.03 | 417 | 0.52 | 0.10 | 5.78 | 2.20 | 0.016 | -735.48 | 30331.75 |
| $b = 5.5$ | 0.78 | 0.94 | 0.02 | 410 | 0.64 | 0.13 | 5.50 | 1.79 | 0.013 | -742.18 | 30599.67 |
| *l = 0.001* | 0.82 | 0.98 | 0.01 | 400 | 0.48 | 0.10 | 4.77 | 2.22 | 0.001 | -730.66 | 30139.17 |
| *c* = 0; *l = 0.001* | 0.79 | 0.94 | 0.00 | 397 | 0.51 | 0.10 | 4.52 | 2.26 | 0.001 | -732.66 | 30104.74 |
| *c* = 0; *l = 0.001; a* = 0.1 | 0.77 | 0.94 | 0.00 | 403 | 0.52 | 0.10 | 5.37 | 2.13 | 0.001 | -742.42 | 30381.13 |
| **Replay Task - no-bound** | | | | | | | | | | | |
| **Model** | $\sigma$ | $\alpha$ | $c$ | $\mu_U$ | $\sigma_U^2$ | $a$ | $b$ | $\lambda$ | $l$ | **LLH** | **ΣBIC** |
| Full | 0.47 | 0.90 | 0.05 | ~ | ~ | ~ | ~ | ~ | 0.012 | -81.13 | 3701.44 |
| $\alpha = 1$ | 0.56 | 1.00 | 0.10 | ~ | ~ | ~ | ~ | ~ | 0.012 | -92.21 | 4030.55 |
| $c = 0$ | 0.48 | 0.90 | 0.00 | ~ | ~ | ~ | ~ | ~ | 0.009 | -82.73 | 3651.38 |
| *l = 0.001* | 0.50 | 0.91 | 0.06 | ~ | ~ | ~ | ~ | ~ | 0.001 | -82.05 | 3624.39 |
| *c* = 0; *l = 0.001* | 0.51 | 0.90 | 0.00 | ~ | ~ | ~ | ~ | ~ | 0.001 | -83.64 | 3573.67 |
| **Replay task - bound** | | | | | | | | | | | |
| **Model** | $\sigma$ | $\alpha$ | $c$ | $\mu_U$ | $\sigma_U^2$ | $a$ | $b$ | $\lambda$ | $l$ | **LLH** | **ΣBIC** |
| Full | 0.44 | 0.87 | 0.10 | ~ | ~ | 0.17 | 8.68 | 11.71 | 0.012 | -79.81 | 3991.09 |
| *c* = 0; *l = 0.001* | 0.48 | 0.88 | 0.00 | ~ | ~ | 0.13 | 8.58 | 15.55 | 0.001 | -82.22 | 3859.24 |
| *c* = 0; *l = 0.001; a* = 0.1 | 0.48 | 0.88 | 0.00 | | | 0.10 | 8.91 | 15.88 | 0.001 | -82.38 | 3751.55 |

***Table S1. Average parameter values.*** *Table shows the average values and the sum of BIC across participants. The large difference in the average loglikelihood (LLH) across tasks is due to the fact the Free task model was fit to both when and what observers responded, whereas in the Replay task only the response was fit. Red values show the fixed parameters. Colour code of the BIC column corresponds to the goodness of fit (the greener the better).*

To compare the Bound and No-bound models in Stage 2. we used five-fold cross validation. The No-bound model had two free parameters: $\alpha$ (temporal bias) and $\sigma$ (inference noise), which were fit to the Same and Less conditions of the Replay task, but tested across all conditions. The Bound model had three free parameters to describe the bound, with the inference noise and temporal bias parameters fixed to those fit to the Same and Less conditions only. In this way, the no-bound model must account for the lack of increased performance in the More condition with the suboptimalities present in the Same and Less conditions, whilst the bound model can limit performance in the More condition in particular by stopping further evidence accumulation. The results of this analysis are presented in the manuscript: the bound significantly improved the fit, mean relative increase in model log-likelihood = 0.048, bootstrapped $p$ = 0.001, **Figure 2c** in the main text.

Of additional interest is the pattern of parameters fit to each condition separately, when the model attempts to explain behaviour without a bound. There was little difference in parameters fit to the Same and Less conditions (mean $\sigma_S = 0.48$, $\sigma_L = 0.44$, $Z(19)$ = -1.46, $p$ = 0.15; $\alpha_S = 0.86$, $\alpha_L = 0.78$, $Z(19)$ = 1.38, $p$ = 0.17). The inference noise fit to the More condition significantly increased from the Less condition ($\sigma_M = 0.55$, $Z(19)$ = -2.61, $p_{bonf*4}$ = 0.036), but there was significantly reduced temporal integration bias ($\alpha_M = 0.93$, $Z(19)$ = -2.50, $p_{bonf*4}$ = 0. 0496) suggesting observers' performance was worse than predicted by the Same and Less conditions, and they were putting less weight on the more recent evidence. These differences in parameters are consistent with the model trying to mimic bounded evidence accumulation without a bound, providing additional support for the comparison described above.

Stage 3. of the model procedure was to account for the confidence ratings. We compared three processing architectures that span the space from single-channel to dual-channel (Maniscalco and Lau, 2016). We took as the null hypothesis a serial processing (single-channel) architecture in which the confidence ratings (Type-II decisions) can be described by the exact same evidence as used to make the perceptual (Type-I) decision. A weaker version of this null hypothesis is that the same suboptimal inference process is used for both perception and confidence, but that the observer can commit to their perceptual decision whilst continuing to monitor additional evidence for evaluating their confidence (a schematic of these processes is shown in **Figure S1a**). The average parameter values are shown in **Table S2**, labelled 'Serial' and 'Serial continued' respectively. Note the substantial increase in inference noise ($\sigma$) and reduction in temporal bias ($\alpha$ is closer to 1) when attempting to describe both the perceptual decision and the confidence rating compared to only the perceptual decision (**Table S1**, Replay task – bound, model $c$ = 0; $l$ = *0.001*). This is indicative of the difficulty of describing both perception and confidence with the same suboptimalities.

At the other extreme is the parallel processing (dual-channel) architecture, in which perception and confidence are computed by independent resources, based on the same sensory input (**Figure S1b**, labelled 'Parallel' in **Table S2**). This is the most computationally expensive description, and provided a lack of parsimony that was only surpassed by a model that attempted to describe confidence ratings with only the inference noise evident from the perceptual decisions.

The intermediate models in this architectural space are the partial dissociation models (**Figure S1c**), which suggest that confidence inherits the same noisy perceptual evidence as the perceptual decision, but may incur some independent suboptimalities. We compared four versions of these models: same $\sigma$ (no additional inference noise); accumulation noise (additional inference noise with each sample of evidence); read-out noise (one additional sample of noise before the confidence response); and same $\alpha$ (the temporal bias affecting the confidence accumulation is the same as that affecting the perceptual accumulation).

In all cases the models were fit to minimise the negative loglikelihood of both perceptual and confidence decisions. The model comparison overwhelmingly favoured the partial dissociation models, and of these, the best description was offered by a model with an independent temporal bias on the confidence evidence accumulation, and additional noise at the read-out stage. We caution against interpreting this result as meaning that there is no additional accumulation noise in the processing of confidence evidence, whilst the models are very similar, it is possible that the read-out noise in this case can additionally capture some noise in setting and maintaining bounds for assigning a rating to the confidence evidence.

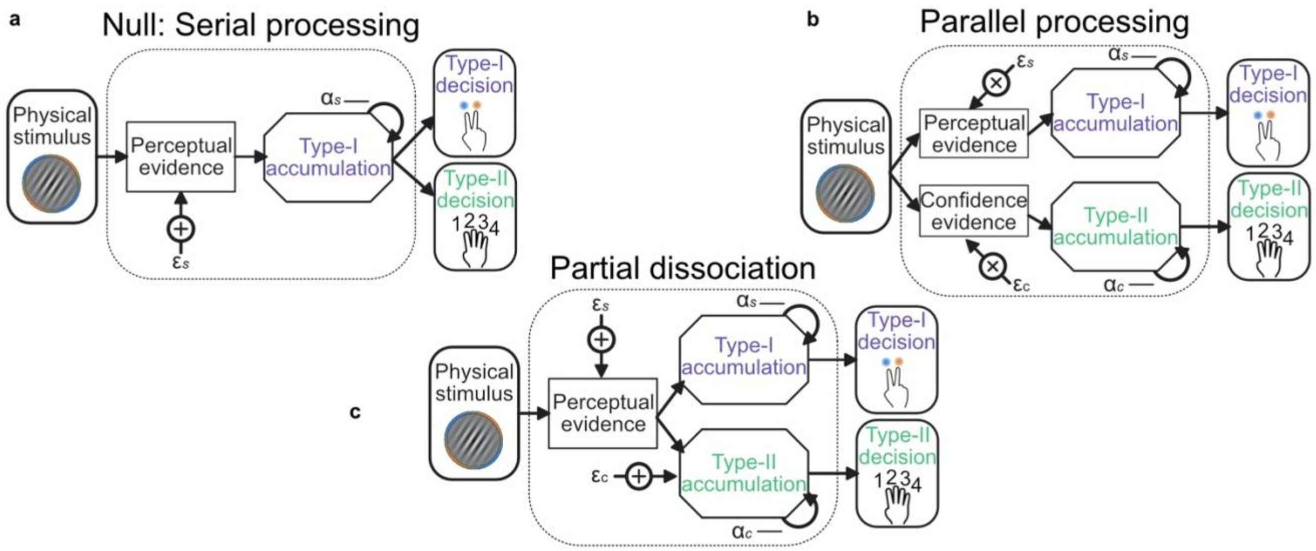


***Figure S1. Schematic of possible relationships between perceptual (Type-I) and confidence (Type-II) evidence accumulation. a)*** *Same evidence accumulation processes: Type-I (perceptual) and Type-II (confidence) decisions are different responses to the same evidence: each sample of perceptual evidence is disrupted by a sample of sensory noise ($\varepsilon_S$) drawn from a zero-mean Gaussian with standard deviation $\sigma$, and accumulated with a temporal bias described by $\alpha_S$.* ***b)*** *Parallel processing: Type-I and Type-II decisions rely on entirely separate processing of the same physical stimulus: the confidence decision also incurs noise and temporal integration bias (with subscript c), but these may vary independently of the perceptual processing suboptimalities (subscript s).* ***c)*** *Partial dissociation: Type-I and Type-II decisions rely on partially dissociable accumulation of the same evidence.*

| Model | $\sigma$ | $\alpha$ | $a$ | $b$ | $\lambda$ | $a_c$ | $b_c$ | $\lambda_{c1}$ | $\lambda_{c2}$ | $\lambda_{c3}$ | LLH | ΣBIC |
|---|---|---|---|---|---|---|---|---|---|---|---|---|
| Serial | 0.73 | 0.92 | 0.10 | 12.74 | 17.07 | 0.07 | 0.64 | 1.28 | 6.81 | 31.38 | -428.36 | 18275.28 |
| Serial continued | 0.67 | 0.91 | 0.13 | 9.60 | 17.98 | 0.06 | 0.53 | 0.66 | 7.08 | 30.41 | -424.88 | 18135.83 |
| Parallel | 0.76 | 0.90 | ~ | ~ | ~ | 0.01 | 0.58 | 0.18 | 7.51 | 30.68 | -437.25 | 18288.50 |
| Partial - same sigma | 0.00 | 0.89 | ~ | ~ | ~ | 0.06 | 0.47 | 1.03 | 6.77 | 25.92 | -446.41 | 18540.68 |
| Partial - accumulation noise | 0.45 | 0.91 | ~ | ~ | ~ | 0.03 | 0.58 | 0.50 | 7.71 | 31.03 | -421.59 | 17662.25 |
| Partial - read-out noise | 0.12 | 0.90 | ~ | ~ | ~ | 0.02 | 0.52 | 1.85 | 8.63 | 37.39 | -417.94 | 17516.29 |
| Partial - same alpha | 0.12 | 0.88 | ~ | ~ | ~ | 0.02 | 0.52 | 0.98 | 8.22 | 35.16 | -423.02 | 17605.29 |

***Table S2. Average parameter values for perceptual and confidence behaviour.*** *Bound parameters with subscript c describe the criteria for confidence ratings, which take the same form as the perceptual decision bound. They have the same minimum and scale, but different rates of decline, such that* $\lambda_{c1}$ *determines the upper bound on a confidence rating of 1, and the lower bound on a rating of 2. Apart from the 'Serial' and 'Serial continued' models, parameters for perceptual decisions were fixed to those fit in the winning perceptual decision model and the listed parameters affect only the confidence evidence accumulation.*

The model comparison of Stage 3. just described mainly assumed continued, unbounded accumulation of confidence evidence (with the exception of the strictly serial processing architecture). Stage 4. was to formally compare bounded and unbounded accumulation for confidence evaluations in the same manner as with the perceptual decisions. This time, two versions of the bound were compared: the same bound as perceptual evidence accumulation (the participant could close their eyes after committing to their perceptual decisions and their responses would not change); or an independent bound (the participant can continue to accumulate evidence for confidence decisions after the committing to the perceptual decision, but will eventually stop). As reported in the manuscript, neither bound improved the fit, if anything, adding the bound decreased the log-likelihood of the model (same bound: relative improvement with bound = -0.007, bootstrapped $p$ = 0.11, uncorrected; independent bound: relative improvement = -0.014, $p$ = 0.022, Bonferroni corrected for two comparisons; **Figure 2c,** in the main text). This reflects the fact that even a very high bound affects the shape of the accumulation trace, which will harm the fit when behaviour is not affected by a bound.

In summary, this computational modelling procedure suggests a partial dissociation in the processing for perception and confidence. In the Replay task, perceptual decisions were best described by bounded evidence accumulation, enabling observers to commit to decisions before the sequence of presented samples finishes. The confidence ratings required additional noise and reduced temporal integration bias compared to the suboptimalities affected the perceptual decisions. These differences were best described by the partial dissociation architecture where confidence received the same noise samples of evidence as the perceptual decision, though they are accumulated differently. In addition, model comparison suggested confidence evidence accumulation continued to the end of the sequence, even in cases of premature commitment to the perceptual decision. The results of these comparisons replicate the results of Balsdon et al. (2020), with the exception of the confidence noise comparison: here we find evidence in favour of read-out noise, whereas the previous analysis found the models indistinguishable.

### Supplementary Note 2

##### Model Simulation

The computational model comparison suggested a partial dissociation in the evidence used to make perceptual decisions and confidence evaluations. We compared the evidence underlying the observers' perceptual decisions and confidence ratings by simulating the winning computational model. For each trial, 10,000 samples of noise per decision update were randomly sampled from the Gaussian distribution describing the observer's inference noise. These were combined to give 10,000 simulated evidence traces per trial. The first 1,000 simulated evidence traces that agreed with the observer's response on that trial were taken to measure the median evidence trace (or, the process was repeated until 1,000 adequate simulated evidence traces were drawn, up to 100 repeats). **Figure S2a** demonstrates this process for one example trial of one observer. For the perceptual evidence (**Figure S2a**, left) simulated evidence traces that agreed with the observer's response are those that reach the respective decision bound before the opposing decision bound, or reach no bound but show evidence in favour of the response by the final sample. It was assumed that once the evidence reaches the bound, that evidence is maintained until the response. For the confidence evaluation (in the example, a confidence rating of 3), the final evidence had to be between the confidence rating bounds to agree with the observer's confidence decision (after the final sample of additional noise – which is why a few samples in **Figure S2a**, right, exceed the bounds). The median evidence was compared to the ideal evidence (green lines of **Figure S2a**).

The estimated inference error (used in **Supplementary Note 7**) scaled the difference between the median consistent evidence and the ideal evidence by the probability of the response given all samples, to estimate the relative deviation of the observers' internal evidence from the optimal observer's evidence. This estimate of the error is quite imprecise: the median trace tends to be quite close to the ideal, even though any one of the traces (which reflect much larger error) could have described the internal evidence of the observer. **Figure S2b** shows the predicted final accumulated evidence for the perceptual (Type-I) compared to the confidence (Type-II) decision for the same example observer. The evidence is strongly correlated but there are substantial deviations, because of the additional noise, different temporal bias, and continued accumulation for the confidence decision, especially in the More condition (light blue). The example observer is a more extreme case because of the relatively strong bound on perceptual evidence accumulation. The (Fisher transformed) correlation for each observer is shown in **Figure S2c**. For many observers there are substantial differences between the median simulated evidence consistent with the perceptual and confidence responses, meaning the simulated evidence could be useful in distinguishing representations important for perception vs. confidence.

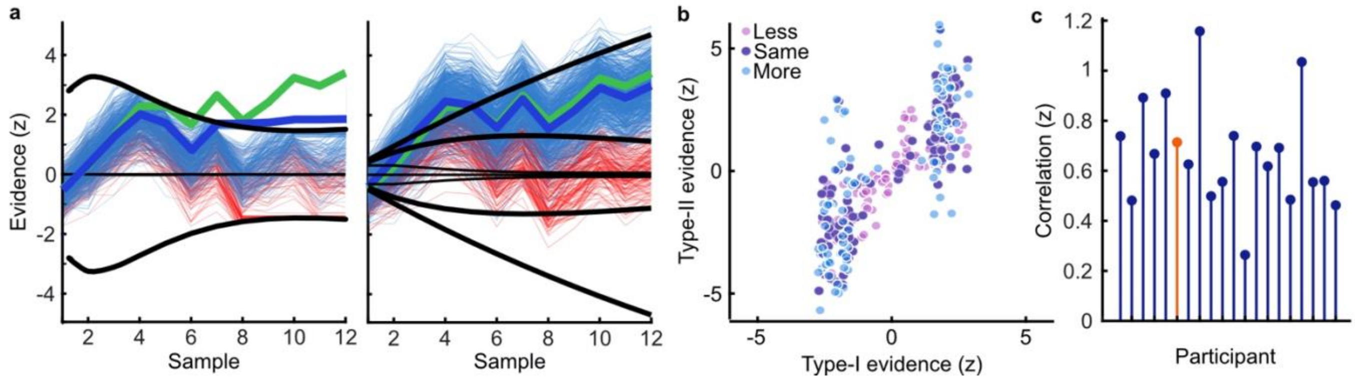


***Figure S2. Model simulation of accumulated evidence for perceptual and confidence decisions. a)*** *Example trial from one observer showing simulated evidence traces agreeing with the observer's response (blue) and a sample of example traces which did not agree (red). The perceptual decision is shown on the left. An evidence trace was taken to agree with the observer's decision if the corresponding bound was reached prior to the opposing bound, or if no bound was reached but the final accumulated evidence was in favour of the chosen option. The median evidence trace (thick blue line) was calculated assuming the evidence that reached the bound early was maintained until the response was entered. For the confidence rating (right) we compared the median evidence from traces where the final accumulator (plus one additional sample of noise) agreed with the observer's confidence rating. We examined the difference from the ideal accumulated evidence (thick green line) relative to the likelihood of the observers' rating given all simulated evidence traces.* ***b)*** *Median final simulated accumulated evidence for the perceptual decision (abscissa), and the confidence decision (ordinate) for all trials of the example observer, colours indicate the condition.* ***c)*** *Correlation (Fisher transformed z) between perceptual and confidence evidence for each observer. The example observer is highlighted in orange.*

### Supplementary Note 3

#### Confidence behaviour

Proportion correct increased with increasing confidence, reflecting the observers' ability to use their confidence ratings to discriminate correct from incorrect responses (**Figure S3a**). Observers appeared to be monitoring the decision evidence to make their confidence ratings, as opposed to some proxy for confidence such as the number of samples they were shown (**Figures S3b** and **S3c**).

We required a single-trial measure of confidence precision for identifying the key neural processes underlying the computation of confidence. To do so, we compared observers' responses to an optimal observer. The optimal observer perfectly accumulates all presented evidence and assigns ratings to equally partition the evidence for their perceptual decision. To simplify, we split trials by the median evidence for the chosen category, where the optimal observer gives a high confidence rating (3 or 4) to those trials with greater than the median evidence, and a low confidence rating (1 or 2) to those with less than the median evidence. We labelled trials as 'suboptimal confidence' when the observer's confidence response disagreed with the response of this optimal observer. This trial labelling is demonstrated for two example observers in **Figure S3d**. We reasoned that on suboptimal confidence trials the internal evidence of the human observer was less likely to be close to the optimal presented evidence, and the neural representation of the optimal presented evidence should be less precise in neural circuits that actually represent this suboptimal confidence evidence. That this measure of confidence precision does capture the suboptimalities in confidence evaluation is confirmed by the significant increase in model estimated confidence error on suboptimal confidence trials (Wilcoxon sign rank test: $Z(19) = 3.85$, $p < 0.001$; **Figure S3e**).

In this way, observers' confidence is assessed relative to a "super-ideal" observer, who has perfect access to the presented evidence (Mamassian and de Gardelle, 2021). Theoretically, observers' confidence should be assessed relative to the internal evidence for their perceptual decision, that is, relative to the evidence based on suboptimal inference (afflicted by noise and temporal integration biases). However, the single-trial estimates of the internal evidence for perceptual decisions, based on model simulations, were relatively imprecise (see **Supplementary Note 2**), and could also introduce systematic errors from the model assumptions, making this estimate of the internal evidence unappealing for the purpose of assessing confidence. Moreover, the goal of this measure was to compare observers' confidence ratings to the neural representation of the accumulated evidence, which was also assessed relative to the optimal evidence. We therefore chose to assess confidence ratings relative to the optimal observer in the same way that neural responses were assessed relative to optimal, though this ignores the fact that some suboptimality is actually inherited from perceptual decision processes.

A second important consideration with this measure is that it is affected by confidence bias. There are three types of biases that could affect confidence ratings: first, a response bias to enter a certain response irrespective of the evidence; second, a miscalibration bias such that ratings mean different things to different observers (the same value of evidence will be given a rating of 4 for one observer and 3 for another, for example); third, a miscalling bias such that perceptual evidence is relatively exaggerated or diminished in the assessment of confidence. All these biases mean that the same internal perceptual evidence could result in systematically different confidence ratings across observers, and observers could report on average higher or lower confidence despite similar perceptual performance and precision in representing the internal evidence for evaluating their confidence.

Taking an average proportion of suboptimal confidence ratings and comparing across observers would result in observers of similar ability having different scores simply because of biases in how they implement the confidence rating responses: greater biases will increase average proportion suboptimal. Importantly, this single-trial measure of confidence was not used for this purpose. Rather, it was compared to neural activity during the process of accumulating evidence for the perceptual decision and confidence evaluation. We expect that biases that are not of interest for the computation of confidence (in particular, response bias and miscalibration bias) are incorporated at a later stage, when the confidence evaluation is converted into a rating for executing the response. The biases will only reduce the sensitivity with which a trial labelled as suboptimal truly reflects internal evidence that differs from optimal, reducing our ability to identify neural processes underlying confidence computation. This is simulated in **Figure S3f**, where a relative bias is introduced by assessing human confidence ratings to a biased optimal observer (who responds on 65% of trials with high confidence – making the human observers relatively more liberal, or 35% high confidence – making the human observers more conservative). The general trend for the difference between confidence ratings that match the (biased) optimal observer and those that are suboptimal remains the same, though the bias reduces the difference.

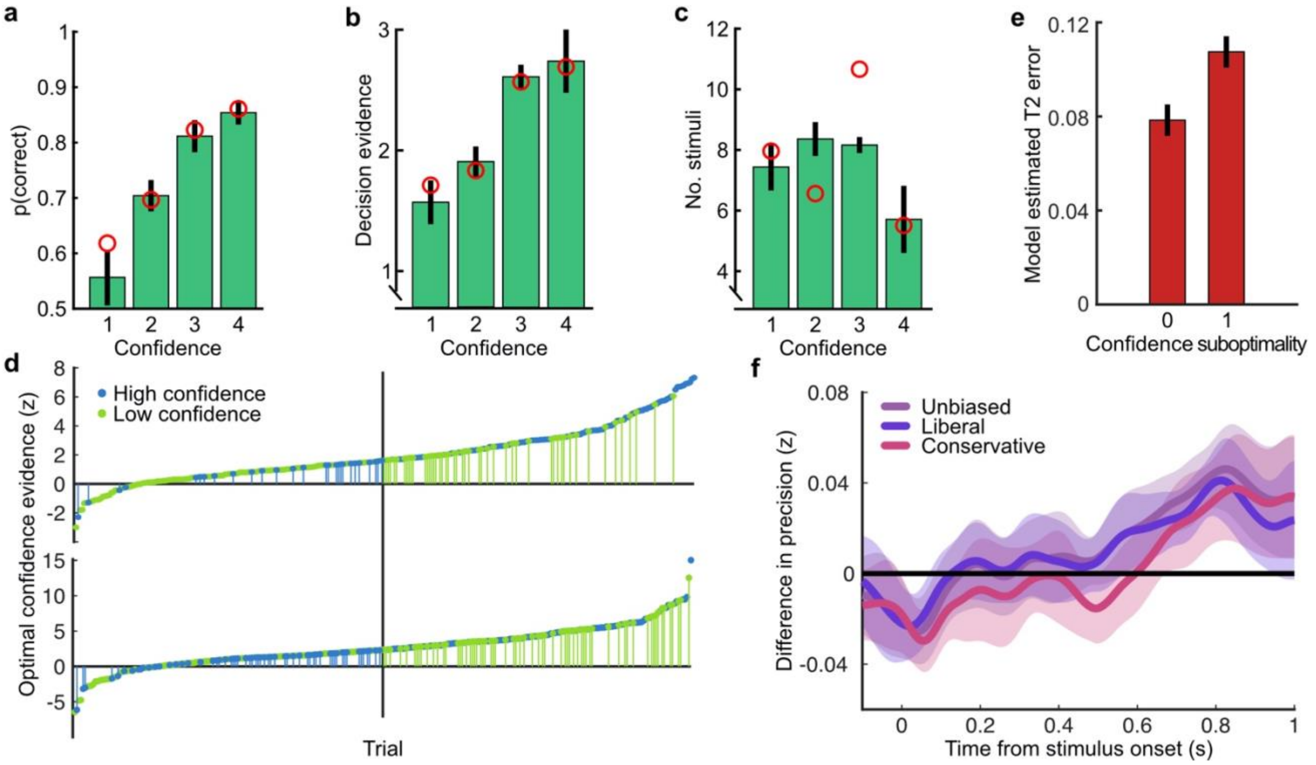


***Figure S3. Confidence behaviour*****.** ***a)*** *proportion correct (in the perceptual decision) by confidence rating.* ***b)*** *Decision evidence (based on the presented samples) by confidence rating.* ***c)*** *Number of samples presented by confidence rating. In all plots, error bars show 95% within-subject confidence intervals. Red circles show the predictions of the best fitting confidence model (**Supplementary Note 1**).* ***d)*** *Confidence responses of two observers (top and bottom panels) on all trials sorted by the confidence evidence of the optimal observer. The median confidence evidence (shown by a black vertical line) defines an optimal confidence observer whose confidence above this median are rated high. Observers' high confidence ratings are shown in blue and low confidence ratings in green. Suboptimal confidence ratings, where human and optimal confidence observers do not match, are indicated with small vertical segments (green for Type-II misses and blue for Type-II false alarms). Negative confidence evidence corresponds to incorrect perceptual decisions. The observer shown on top clearly has fewer suboptimal responses compared with the observer below, and the frequency of suboptimal responses decreases further from the median.* ***e)*** *Model estimated confidence error by confidence rating suboptimality (0 = the observer's confidence rating was the same as the optimal observer, 1 = suboptimal confidence rating).* ***f)*** *The effect of response bias on the analysis of suboptimal confidence in the EEG representation of accumulated evidence. Observers' confidence ratings were compared to an unbiased optimal observer (purple), and two biased (but otherwise optimal) observers, who respond with high confidence on 35% and 65% of trials (making the human observers relatively more liberal and conservative with their response strategy in comparison). Thick lines show the within-subject difference in precision (Fisher transformed*

*correlation) between trials where the human observers' confidence ratings correspond to the (un/biased) optimal observer and suboptimal confidence ratings. Shaded regions show the 95% between-subject confidence intervals on the difference.*

### Supplementary Note 4

**Classical EEG analyses**

To link back with the previous literature, we present here two more classical EEG analysis approaches, examining the modulations of EEG amplitude around the time of the response. In **Figure S4a,** we show the Lateralised Readiness Potential (LRP; difference in microvolts between the average of electrodes [C1, C3], and [C2, C4], signed by response hand; Deecke et al., 1976). The data are unfiltered with the exception of the pre-processing, and baselined using the 100 ms before the onset of the first stimulus of each trial. There was a significant difference in the LRP between the Less and More conditions of the Replay task from just after the response (the first cluster from 32 to 196 ms; $t_{\mathrm{ave}}(19) = -3.57$, $p_{cluster} < 0.002$, Figure S4a, top). There were also differences based on perceptual decision accuracy (from -84 ms to 652 ms around the response, with the largest difference just after the response, $t_{\mathrm{ave}}(19) = 2.81$, $p_{cluster} < 0.002$; **Figure S4a**, middle). There was no significant difference in the LRP between trials with high confidence (ratings of 3 and 4) and low confidence (ratings of 1 and 2; **Figure S4a**, bottom).

We also computed the Central Parietal Positivity (CPP; O'Connell et al., 2012) which has previously been shown to reflect perceptual evidence accumulation. We followed the methods presented in Kelly and O'Connell, 2013: data were lowpass filtered at 45 Hz with no highpass filter, and converted to current source density (Kayser and Tenke, 2006). As with the LRP, a baseline was taken from the 100 ms before the onset of the first stimulus of each trial. The slopes of the CPP (a linear fit from -500 to -50 ms) showed no significant differences across all conditions ($F(1,19) = 2.15$, $p = 0.14$). We observed a significantly greater slope for correct compared to incorrect decisions ($t(19) = -2.86$, $p = 0.01$), and an even greater difference between high and low confidence trials ($t(19) = -3.24$, $p = 0.004$). This is consistent with the literature suggesting the CPP traces the internal evidence for the perceptual decision, however it is difficult to disambiguate how this signal may differentially contribute to perceptual decisions and confidence evaluations.

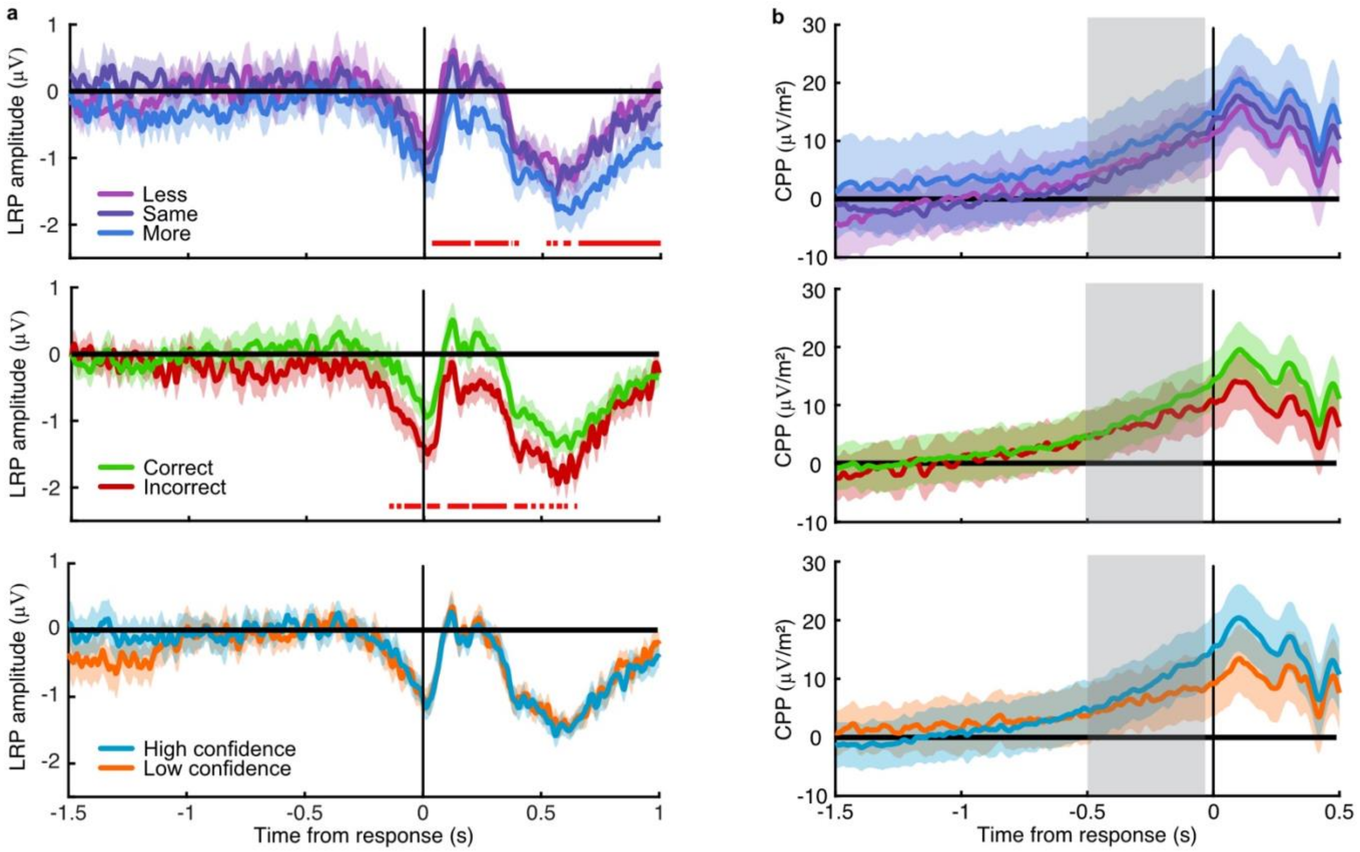


*Figure S4. Amplitude modulations with task variables.* ***a)*** *The Laterised readiness potential by condition (top), perceptual decision accuracy (middle) and reported confidence (bottom). Horizontal red lines mark significant differences in amplitude.* ***b)*** *Central Parietal Positivity, with the same comparisons. Shaded regions show 95% within subject confidence intervals, and the region of slope comparison for the CPP is highlighted in grey.*

### Supplementary Note 5

##### Response classification

A linear discriminant analysis was used to classify the perceptual decision response based on the spectral power of band-limited EEG signals in epochs locked to the time of the response. The spectral power across frequency tapers from 1 to 64 Hz with 25% spectral smoothing was resolved using wavelet convolution implemented in FieldTrip (Oostenveld et al., 2011). The epochs were then clipped at -3 to 1 s around the time of entering the perceptual decision response. We first trained and tested at each frequency taper at each time point in the Free task (**Figure S5a**). Classifier performance was measured as the area under the curve (AUC). The power in frequency bands between 8 and 32 Hz yielded the most accurate classification performance. The difference in the average power across these frequency bands between -0.5 and 0.5 seconds around the time of the response for right- and left-handed responses showed a clear lateralisation over central and parietal electrodes (**Figure S5b**). Training and testing at each time point in each condition of the Replay task showed a similar pattern to the Free task, with reliable classifier performance from around -0.5 to 0.5 seconds around the response (**Figure S5c**). Training and testing within each condition of the Replay task resulted in a larger between-subject error, likely because there are only 100 trials per condition. In the main text, we present a cross-classification analysis where the classifier is trained on the Free task, and tested on each condition in the Replay task, which more directly examines when the signals relevant for entering a response (based on the Free task) emerge during the lead up to the response in each condition of the Replay task.

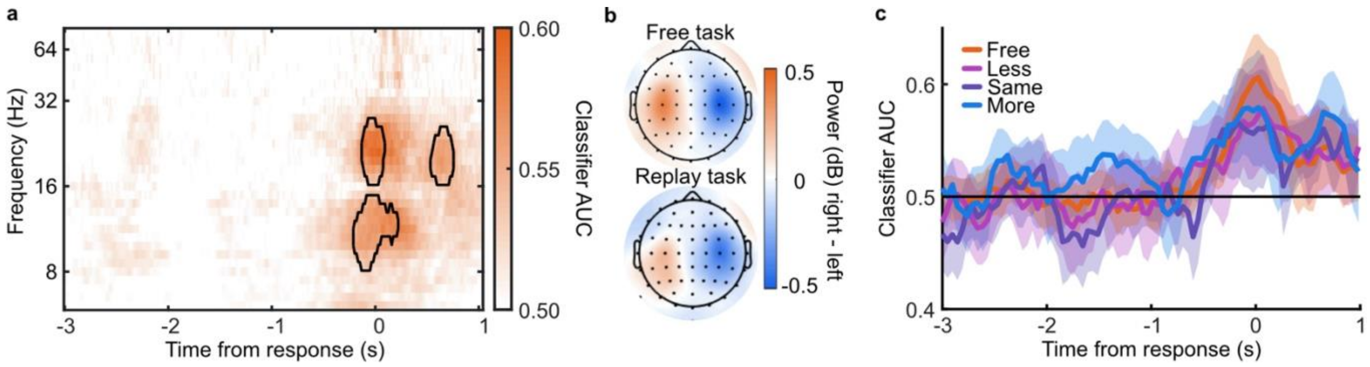


***Figure S5. Response Classification analysis. a)*** *Classifier AUC training and testing at each time point (abscissa) based on the power (dB) in each frequency band (ordinate). Clusters where average performance is greater than 3.1 standard deviations (99% confidence) from baseline (0.5) are circled in black.* ***b)*** *Scalp map of the difference in power for right- compared to left-handed responses averaged over 8 to 32 Hz and -0.5 to 05 seconds around the response.* ***c)*** *Classifier performance (AUC) training and testing at each time point, in each condition of the Replay task and in the Free task.*

### Supplementary Note 6

##### Encoding variable regression

Linear regression was used to examine the representation of encoding variables in the EEG signals. First, regression weights ($\widehat{W}$) were computed using ridge regression of the encoding variables ($C$, an *n × 1* matrix) on the EEG signals ($D$, an *n × m* matrix, where *m* is the number of EEG signals, and *n*, the number of epochs)

$$\widehat{W} = (D^T D + \lambda I)^{-1} D^T C \qquad \textbf{(4)}$$

The regularisation parameter, $\lambda$, was set to 1, where $I$ is the identity matrix. Weights were computed on 90% of the epochs, and used to predict the encoding variables on the other 10% (10-fold cross validation) simply as: $\hat{C} = D * \widehat{W}$. The precision of the prediction was calculated as the correlation between $\hat{C}$ and $C$, standardised using a Fisher transformation.

Three different encoding variables, $C_\theta$, $C_\ell$, and $C_z$, were examined (**Figure S6a**): the stimulus orientation ($C_\theta = \pi - |\theta_n|$), the momentary decision update ($C_\ell = |\ell_n| = \left| \kappa cos\big(2(\theta_n - \mu_1)\big) - \kappa cos\big(2(\theta_n - \mu_2)\big) \right|$), and the accumulated evidence ($C_z = z_n = \sum_{N=1}^{n} \ell_N$ , signed by the response). These variables are not entirely independent: There is a weak correlation between the stimulus orientation and the momentary decision update ($r$ = 0.03), and a weak correlation between the momentary decision update and the accumulated evidence ($r = 0.09$). In addition, the accumulated evidence is strongly correlated over samples ($r = 0.92$ at n+1, and $r = 0.85$ at n+2). The cross-correlations are shown in **Figure S6c**.

The EEG signals in D were low-pass filtered and decomposed into real and imaginary parts using a Hilbert transform. Regression precision was first calculated using the signals from all electrodes ($m = 128$) separately for each time-point in the stimulus-locked epochs. Initial analysis showed a low-pass cut-off of 8 Hz was appropriate to decrease noise whilst maintaining precision (**Figure S6b**). The previous literature has shown similar results (Salvador et al., 2020).

Temporal generalisation of the representation of encoding variables was tested by computing weights at each time point and testing the predicted encoding variables across time (**Figure S6d**). Though the representation of the momentary decision update is maintained for a relatively longer duration than the representation of stimulus orientation, there is little temporal generalisation, suggesting the representation in the EEG signals evolves over time. This is also the case for the representation of accumulated evidence, however, there are also strong off-diagonals in the temporal generalisation matrix. This is likely because of the strong correlation across consecutive samples (**Figure S6c**).

The precision of the representation of accumulated evidence was compared across the Less and More conditions for the first four and the last four stimuli (**Figure S6e**). As reported in the main text, representation precision was substantially attenuated for the last four stimuli of the More condition. This was not the case for the first four samples, where decoding precision in the More condition was briefly (from 132 to 244 ms) greater than in the Less condition ($t_{ave}(19) = 3.67$, $p_{cluster} < 0.001$).

Given the sustained precision of decoding accumulated evidence over time, and the strong correlation between consecutive samples, it is curious that the measured precision does drop to baseline at the start of the epoch. That the same pattern is found when decoding sample n-1 and sample n+1 based on the epoch at sample n (**Figure S6f**) suggests that the onset of the stimulus is disrupting the ongoing representation (or at least, our ability to measure it). Furthermore, this decrease in performance is not seen in the temporal generalisation matrix, where the off-diagonal is not aligned with the onset of successive samples (due to the jitter in stimulus presentation timing). Comparing precision between groups of epochs where the timing of the subsequent sample is aligned (**Figure S6g**; red 317 ms, green 333 ms, blue 350 ms) suggests there could be an interaction between the timing of ongoing updates and the precision of the representation of the accumulated evidence (but not the momentary decision update). This could be of interest for future research.

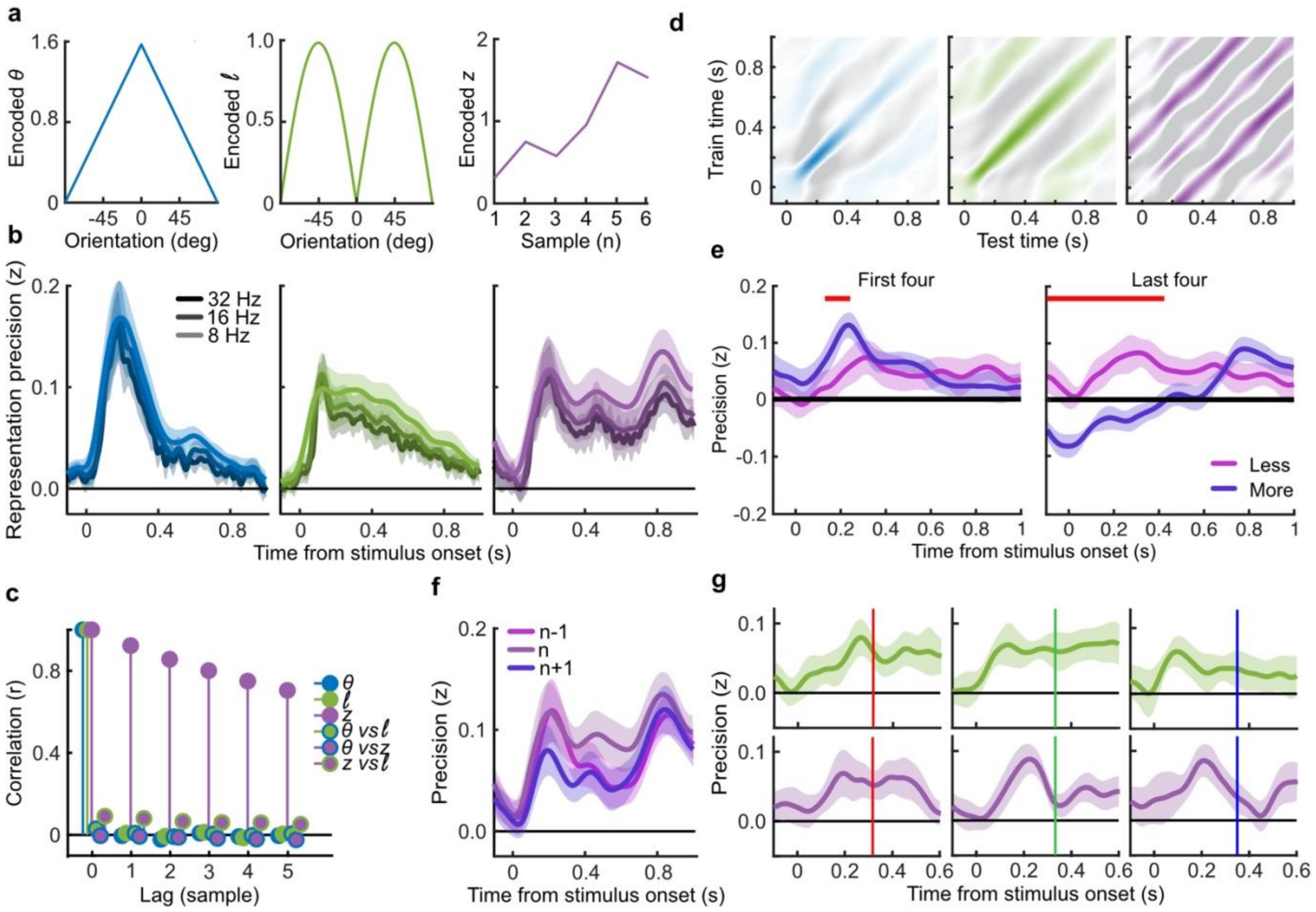


***Figure S6. Encoding variable regression. a)*** *Encoded variables used to regress EEG signals. The encoded orientation ($C_\theta$, left) and encoded momentary decision update ($C_\ell$, middle) were dependent on the orientation presented to the observer. The encoded accumulated evidence ($C_z$) varied over all presented orientations in a trial, the figure on the right shows only one example.* ***b)*** *Representation precision of encoding variables using different low-pass filters.* ***c)*** *Cross correlation between encoding variables over consecutive samples.* ***d)*** *Temporal generalisation of representations: the regression weights were calculated on EEG signals at each time point and precision was tested across time. Colour scales are relative to the maximal precision, with zero precision in white and negative in grey (a sign flip of the regression weights).* ***e)*** *Representation precision of the accumulated evidence for the first (left) and last (right) four stimuli of the Less and More conditions. Shaded error bars show the 95% within subject confidence intervals, red horizontal bars mark cluster corrected significant differences between conditions.* ***f)*** *Representation precision of the previous (n-1), current (n) and future (n+1) accumulated evidence, based on the EEG signals locked to the current epoch.* ***g)*** *Representation precision of the momentary decision update (top) and the accumulated evidence (bottom) for epochs separated by the timing of the subsequent stimulus, shown in coloured bars (317 ms, red, left; 333 ms, green, middle; and 350 ms blue, right).*

### Supplementary Note 7

##### Cluster modelling

Cluster modelling was used to isolate contiguous signals in space (electrode location) and time, where the precision of the representation of optimal accumulated evidence systematically varied how closely the internal representation of evidence matched optimal, based on whether behavioural responses matched the optimal observer. We assume that responses that do not match the optimal observer are based on evidence that deviates further from the optimal evidence. Neural signals that reflect the internal evidence of the observer should also deviate further from the optimal evidence used in the regression on these trials, and be closer to the optimal evidence on trials where the observers response matched the optimal observer. Clusters were isolated using a multivariate Bayesian scan statistic (Neill, 2011; Neill, 2019). This statistic was calculated based on the loglikelihood ratio of the alternative hypothesis (that representation precision depends on the internal evidence of the observer) against the null hypothesis (that any difference in representation precision is due to measurement noise alone, which is independent across epochs). It is assumed that the neural signals reflect the input (cumulative presented evidence) with added measurement noise ($N_m$) and, when the neural signals are relevant for behaviour, inference noise ($N_i$) that reflects the deviations from the optimal evidence in the internal representation of the observer

$$Y_{out} = Y_{in} + N_i + N_m \quad \textbf{(5)}$$

Where the two sources of noise are assumed to be gaussian distributed ($N(0, \sigma^2)$). The total measured correlation ($r_T$) between $Y_{in}$ and $Y_{out}$ is a function of the additional noise (where $Y_{in}$ is normalised)

$$r_T = \frac{1}{\sqrt{2 + \sigma_i^2 + \sigma_m^2}} \quad \textbf{(6)}$$

When the observer's decision does not match the optimal decision their internal representation of the accumulated evidence is likely to be further from the optimal value, resulting in a weaker correlation between the internal representation and the presented evidence. Therefore, when we split based on behaviour, we expect that on average there is greater inference noise on incorrect trials than correct trials. The correlation over all samples can be described as

$$r_T = \frac{1}{\sqrt{2 + p(I)\sigma_{iI}^2 + p(C)\sigma_{iC}^2 + \sigma_m^2}} \quad \textbf{(7)}$$

where *p(I)* is the observed probability of a decision that does not match the optimal observer, and *p(C),* a decision that corresponds to that of the optimal observer. The null hypothesis is that the neural signal is not relevant for behaviour, specifically, signals on suboptimal trials do not reflect additional inference noise. Any difference in the correlation is due to variance in the measurement noise,

$$H_0\colon\ \sigma_{iI} = \sigma_{iC} = 0 \quad \textbf{(8)}$$

The alternative hypothesis is that the neural signals are relevant for behaviour, reflecting the greater variance from optimal on trials where the observer makes a decision that does not match the optimal decision,

$$H_1\colon\ \sigma_{iI} > \sigma_{iC}\text{ , or }\sigma_{iI}^2 = (\sigma_{iC}^2 - x)\text{ where }x > 0 \quad \textbf{(9)}$$

The difference in the inference noise is limited by the total variance

$$p(I)(\sigma_{iI}^2) + p(C)(\sigma_{iI}^2 + x) = \frac{1}{r_T^2} - 2 - \sigma_m^2 \quad \textbf{(10)}$$

Solving for $\sigma_{iI}^2$ (since $p(C) + p(I) = 1$):

$$\sigma_{iI}^2 = \frac{1}{r_T^2} - 2 - \sigma_m^2 - p(C)x \quad \textbf{(11)}$$

If we consider the correlation between the neural representation and the presented evidence on trials with optimal and non-optimal responses separately (for simplicity, let $R = \frac{1}{r_T^2}$ ),

$$r_I = \frac{1}{\sqrt{R - p(C)x}} \quad \textbf{(12)}$$

$$r_C = \frac{1}{\sqrt{R - p(C)x - x}} \quad \textbf{(13)}$$

Setting a uniform prior on the ratio of inference and measurement noise, results in a linearly descending prior on $x$

$$p(x) = \frac{R - 2 - p(I)x}{\int_0^{(R-2))/p(I)} R - 2 - p(I)x\ dx} \quad \textbf{(14)}$$

We actually measure the difference in the Fischer transform of the correlation

$$z_C - z_I = 0.5\log\left(\frac{(1 + r_c)(1 - r_I)}{(1 - r_c)(1 + r_I)}\right) \quad \textbf{(15)}$$

Since $r_c$ and $r_I$ are independent of the assumed measurement noise, there is one $x$ that corresponds to a measured difference $z_C - z_I$, given the overall correlation $r_T$.

For each participant, for each electrode, at each time-point, the prior on $\sigma_m^2$ for $H_0$ is calculated by permuting the data labels (accurate vs inaccurate behavioural responses). The probability of the data given $H_0$ and $H_1$

are calculated as above and used to compute the loglikelihood ratio

$$LLR = log\left(\frac{p(D|H_1)}{p(D|H_0)}\right) \quad \textbf{(16)}$$

The clusters are identified using the Fast Subset Sums procedure: The loglikelihood ratios are summed across participants, for each electrode and time-point. We then find small clusters by thresholding the log posterior odds ratio

$$POR = LLR + log\left(\frac{p(H_1)}{p(H_0)}\right) \quad \textbf{(17)}$$

where the prior $p(H_1)$ is set to 0.05. The cluster with the largest LLR (summed across electrodes and time points) is then expanded by continuing to add the largest neighbour and the new log prior ($p(H_1) = 0.05/n$), where $n$ is the size of the cluster, whilst the POR remains in favour of $H_1$. This is repeated until all clusters with evidence in favour of $H_1$ have been identified.

### Supplementary Note 8

**Estimating single-sample confidence inference error**

We aimed to examine the neural processes that are important for the representation of the decision evidence for computing confidence. To do so, we explored the source(s) of the activity reflecting the neural representation of the accumulated evidence in the clusters of signals identified as relevant for confidence evaluations. We use the representation from the cluster as an estimate of the internal evidence the observer uses to make their confidence evaluations. The cluster inference error is the absolute difference between the predicted value (on each sample) and the optimal value given the presented evidence. We take this as an estimate of the inference error of the observer at the sample level. This estimate is likely substantially affected by measurement noise, however, we do not expect measurement noise to be systematically driven by a specific source, especially not across subjects. Noise Min and Noise Max epochs were selected by taking the top and bottom quartiles of epochs sorted by this estimate of inference error.

A separate estimate of the inference error was obtained by simulating the computational model (**Figure S7a** shows the process of obtaining these estimates and their mutual reliance on the input stimulus variables and the behavioural output). This computational model estimate also has its drawbacks: It is relatively imprecise, given the large range of errors that are consistent with the observers' behavioural responses (see **Supplementary Note 2**); and is based on the assumptions of the model. By examining these two estimates, we avoid relying on the same set of assumptions throughout the analysis. As reported in the **Results** section, the estimate of the single-sample inference error from the cluster representation was significantly correlated with the single-sample inference error estimated from the computational model of confidence ratings ($t(19) = 5.12$, $p < 0.001$), and this correlation was significantly greater than the error estimated from the model of perceptual decisions alone ($t(19) = 2.62$, $p = 0.017$). This correlation between these estimates suggests that they do tap into the suboptimal inference of the observer.

**Figure S7b** shows the correlation of these estimates of the inference error and different variables related to the stimulus presentation and behaviour, averaged across subjects. We also examined the average absolute effect size of the within subject difference between different variables dividing trials by Noise Min and Noise Max epochs is shown in **Figure S7c**. There was a larger effect on confidence inference error ($d$ = 0.06) than perceptual inference error ($d$ = 0.02), from the model estimate. There were some effects on stimulus variables: a small effect of condition (More vs Less, $d$ = 0.03), a large effect on sample position in the sequence (Noise Min epochs tended to correspond to earlier samples, $d$ = 0.2), and an effect on decision update (Noise Min epochs tended to correspond to smaller momentary decision updates, $d$ = 0.08). The effects on behaviour were largest for confidence accuracy ($d$ = 0.06), with limited effect on perceptual accuracy ($d$ = 0.02) and confidence rating (Noise Min epochs were somewhat more associated with high confidence ratings, $d$ = 0.03).

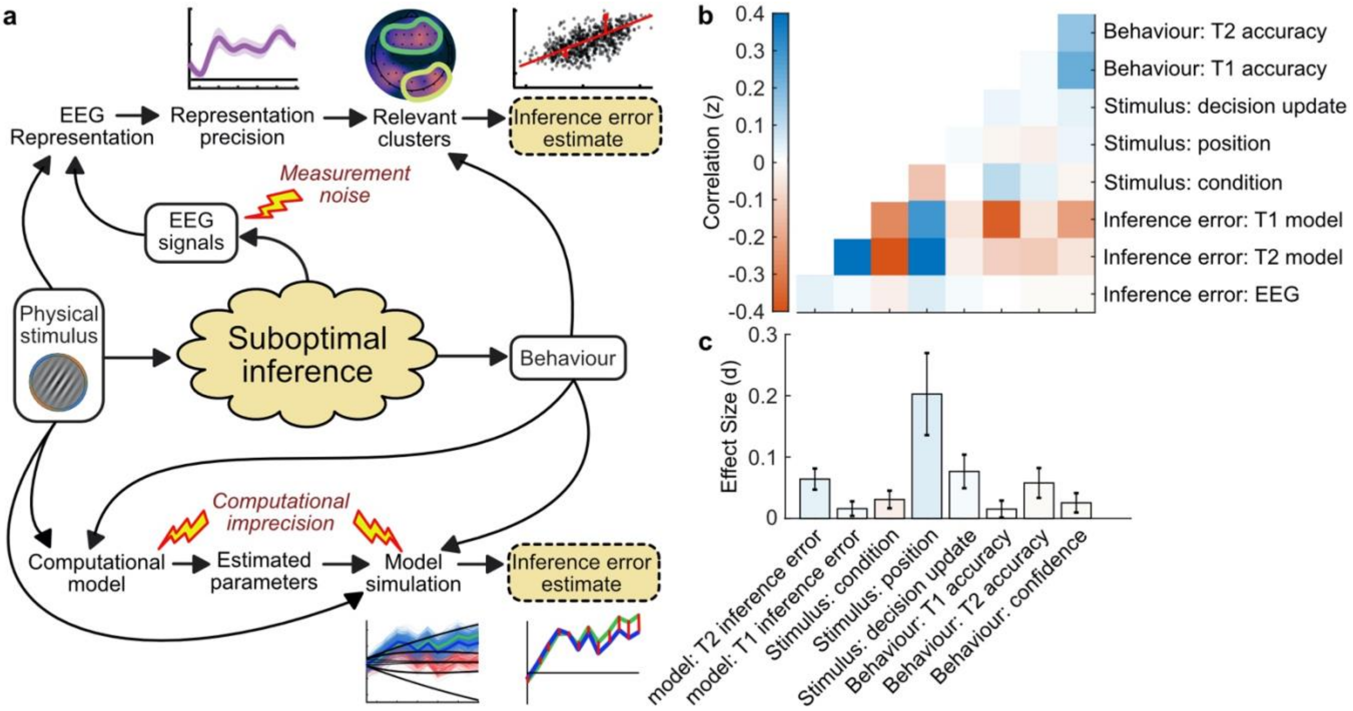


***Figure S7. Estimating inference error. a)*** *Two approaches to estimate inference error. It is assumed the observer's behaviour is based on a suboptimal inference over the physical stimulus. We do not have access to the single-sample inference error, but can estimate it using the measured variables: the physical stimulus properties, the behaviour, and the EEG signals. Two approaches are outlined: The EEG inference error estimate, which relies on the error of the representation of the accumulated evidence, in clusters where the precision of the representation is related to suboptimal behaviour; and the model error, which relies on simulating the processing of the evidence based on the fitted model parameters, and taking the median of simulated traces which concur with the observer's response.* ***b)*** *Correlation between variables measured from behaviour, the stimulus input, and the estimated inference error.* ***c)*** *Effect size on the difference between Noise Min and Noise Max epochs.*

### Supplementary Note 9

**Regions of interest**

Regions of interest were selected based on the previous literature. Specifically, Herding et al. (2019) found subjective evidence to modulate activity in the superior parietal cortex; Gherman and Philiastides (2018) found correlates of confidence encoding in the ventro-medial prefrontal cortex (overlapping with the MindBoggle orbitofrontal cortex coordinates), whilst Graziano et al., (2015) examined ROIs in the anterior cingulate cortex, orbitofrontal cortex, temporal lobe, posterior parietal cortex, and occipital cortex. We chose to use ROIs defined by MindBoggle (Klein et al., 2017) that corresponded to similar regions: lateral occipital cortex, superior parietal cortex, orbitofrontal cortex (combining medial and lateral partitions), rostral middle frontal cortex, and initially the anterior cingulate cortex (combining rostral and caudal partitions; **Figure S8b**). These regions do not necessarily map on to regions of the greatest current density (**Figure S8a** shows the current density over time for the Noise Min epochs). The results of the anterior cingulate cortex were similar to the neighbouring orbitofrontal region, so we decided not to present this in the manuscript for simplicity. We show the results in **Figure S8c**, for left and right hemispheres separately (statistical analyses were performed on the average).

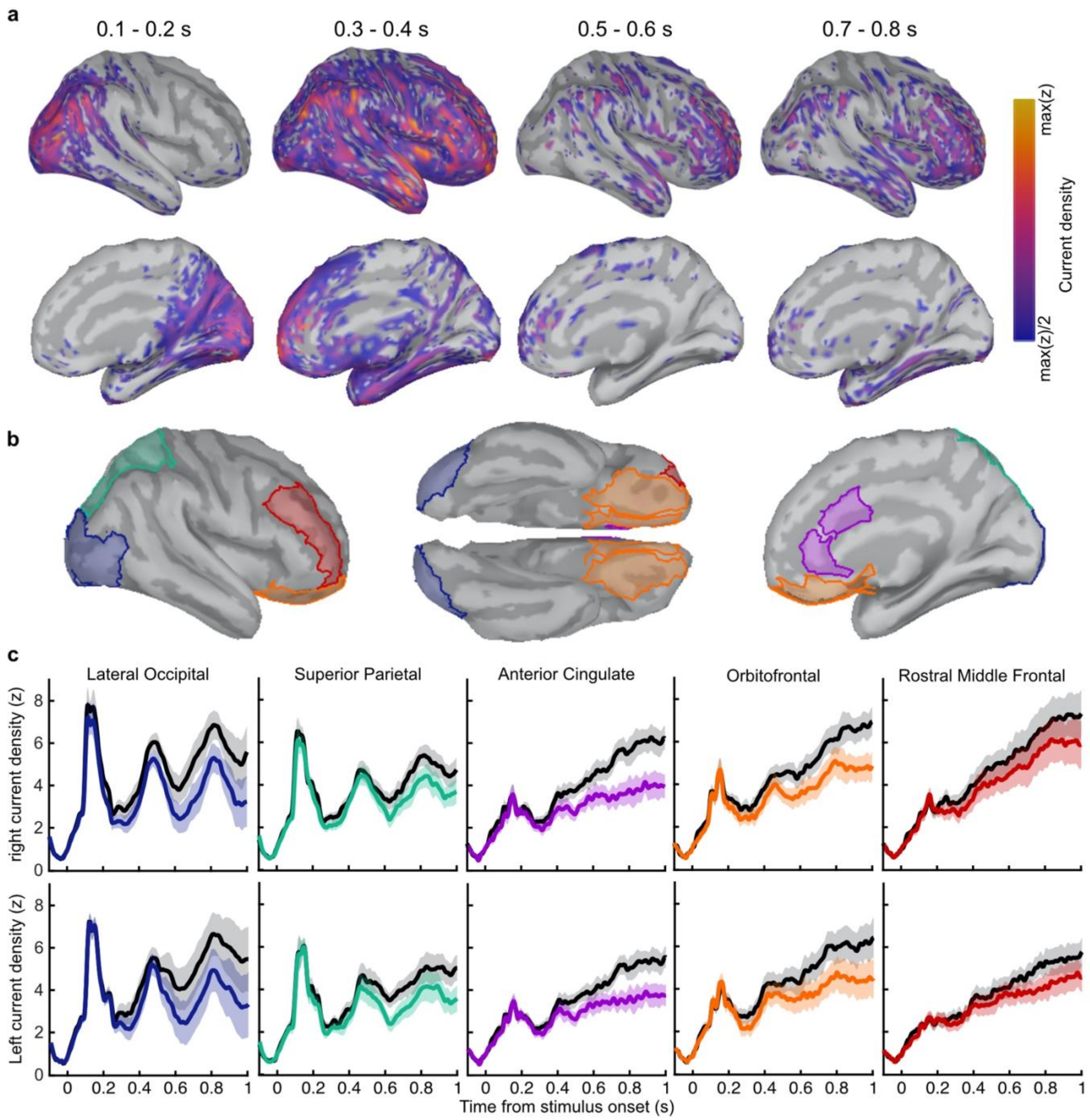


486

487 ***Figure S7. Regions of interest and corresponding current density. a)*** *Average rectified normalised current*
488 *density in Noise Min epochs for the corresponding time windows, filtered above the half-maximum amplitude* ***b)***
489 *Regions of interest based on Mindboggle coordinates.* ***b)*** *Average normalised rectified current density in the*
490 *right (top) and left (bottom) hemispheres. Noise Min epochs are shown coloured, Noise Max in black, with*
491 *shaded regions showing the 95% within-subject confidence interval.*

#### References


Balsdon, T., Wyart, V., & Mamassian, P. Confidence controls perceptual evidence accumulation. *Nature Communications*, 2020; **11**(1), 1-11

Deecke, L., Grozinger, B., & Kornhuber, H. H. Voluntary finger movement in man: Cerebral potentials and theory. *Biological Cybernetics*, 1976; **23**, 99–119.

Kelly, S. P., & O'Connell, R. G. Internal and external influences on the rate of sensory evidence accumulation in the human brain. *Journal of Neuroscience*, 2013; ***33***(50), 19434-19441.

Klein, A., Ghosh, S. S., Bao, F. S., Giard, J., Häme, Y., Stavsky, E., ... & Keshavan, A. Mindboggling morphometry of human brains. *PLoS Computational Biology,* 2017; **13**(2), e1005350.

Mamassian, P., & de Gardelle, V. Modeling perceptual confidence and the confidence forced-choice paradigm. *Psychological Review, 2021.*

Maniscalco, B., & Lau, H. The signal processing architecture underlying subjective reports of sensory awareness. *Neuroscience of Consciousness*, 2016; **1**.

Neill, D. B. Fast Bayesian scan statistics for multivariate event detection and visualization. *Statistics in Medicine*, 2011; **30**(5), 455-469.

Neill, D. B. Bayesian Scan Statistics. In: Glaz J., Koutras M. (eds) *Handbook of Scan Statistics.* 2019*;* Springer, New York, NY.

O'Connell, R. G., Dockree, P. M., & Kelly, S. P. A supramodal accumulation-to-bound signal that determines perceptual decisions in humans. *Nature Neuroscience*, 2012; **15**(12), 1729.

Oostenveld, R., Fries, P., Maris, E., & Schoffelen, J. FieldTrip: open source software for advanced analysis of MEG, EEG, and invasive electrophysiological data. *Computational Intelligence and Neuroscience*, 2011.

Salvador, A., Arnal, L. H., Vinckier, F., Domenech, P., Gaillard, R., & Wyart, V. Premature commitment to uncertain beliefs during human NMDA receptor hypofunction. bioRxiv 2020
